# Anaerobic microbial degradation of protein and lipid macromolecules in subarctic marine sediment

**DOI:** 10.1101/2020.04.27.061291

**Authors:** Claus Pelikan, Kenneth Wasmund, Clemens Glombitza, Bela Hausmann, Craig W. Herbold, Mathias Flieder, Alexander Loy

**Affiliations:** Division of Microbial Ecology, Centre for Microbiology and Environmental Systems Science, University of Vienna, Vienna, Austria; Austrian Polar Research Institute, Vienna, Austria; Center for Geomicrobiology, Department of Biology, Aarhus University, Aarhus, Denmark; ETH Zürich, Department of Environmental Systems Science, Zürich, Switzerland; Joint Microbiome Facility of the Medical University of Vienna and the University of Vienna, Vienna, Austria; Department of Laboratory Medicine, Medical University of Vienna, Vienna, Austria

## Abstract

Microorganisms in marine sediments play major roles in marine biogeochemical cycles by mineralizing substantial quantities of organic matter from decaying cells. Proteins and lipids are abundant components of necromass, yet microorganisms that degrade them remain understudied. Here, we revealed identities, trophic interactions and genomic features of microorganisms that degraded ^13^C-labelled proteins and lipids in cold anoxic microcosms with sulfidic subarctic marine sediment. Supplemented proteins and lipids were rapidly fermented to various volatile fatty acids within five days. DNA-stable isotope probing (SIP) suggested *Psychrilyobacter atlanticus* was an important primary degrader of proteins, and *Psychromonas* members were important primary degraders of both proteins and lipids. Closely related *Psychromonas* populations, as represented by distinct 16S rRNA gene variants, differentially utilized either proteins or lipids. DNA-SIP also showed ^13^C-labeling of various *Deltaproteobacteria* within ten days, indicating trophic transfer of carbon to putative sulfate-reducers. Metagenome-assembled genomes revealed the primary hydrolyzers encoded secreted peptidases or lipases, and enzymes for catabolism of protein or lipid degradation products. *Psychromonas* were prevalent in diverse marine sediments, suggesting they are important players in organic carbon processing *in situ*. Together, this study provides an improved understanding of the metabolic processes and functional partitioning of necromass macromolecules among microorganisms in the seafloor.

## Introduction

The majority of marine sediments that underlie the Earth’s oceans are dominated by microorganisms mediating heterotrophic processes, which are primarily sustained by “pelagic-benthic” coupling [1]. This is driven by a constant supply of organic matter from planktonic organisms that thrive in the overlying water column and settle with particulate aggregates to the seafloor after their death [2]. The amount and composition of organic matter that reaches the seafloor is strongly dependent on water depth, whereby continental shelves and shallow sediments generally receive greater inputs compared to deep sea sediments [3]. In cold polar regions with lower microbial activities in the water column [4], a large fraction of the phytodetritus from spring bloom-like events reaches the underlying sediments [5]. Additionally, organic matter can originate from autochthonous microbial production [6] and from land-derived sources, which can be substantial in areas close to land and rivers, as well as polar regions subjected to melting [2, 7].

Organic matter in marine sediments is composed of an immense diversity of macromolecules, of which a substantial fraction can be enzymatically degraded and utilised as nutrient and energy sources by microorganisms [2]. Proteins and lipids typically constitute 10% and 5-10% of the organic matter found in marine sediments, respectively [8, 9]. Proteins, peptides and amino acids are important sources of nitrogen [10], particularly in sediment habitats with limited inorganic nitrogen sources [11]. Lipids generally consist of an alkyl chain that is ester- or ether-bound to a polar head group such as phospho-glycerol, or a glycerol that is glycosidically bound to a sugar moiety. They are major components of phytoplankton biomass [12, 13], of which the less labile fraction survives degradation in the water column [14] and is degraded by microorganisms in the underlying sediments [15]. Long-chain fatty acids released from lipid hydrolysis are energy-rich compounds [16], and can be expected to be favourable substrates for microorganisms.

Most macromolecules are too large to be directly imported into cells and must therefore be degraded, at least partially, outside the cells [17]. Hydrolysis of macromolecules via the activity of extracellular enzymes is the rate limiting step during organic matter mineralization in heterotrophic marine sediments [17, 18]. Microorganisms utilize a compositionally and functionally large diversity of extracellular hydrolases to facilitate the extracellular breakdown of macromolecules [19]. For example, peptidases in sediments of Arctic fjords in Svalbard had greater substrate ranges for different peptides and much higher activities (tens to hundreds of thousands nmol l^−1^ hr^−1^) than peptidases in the water column [20]. This showed that sediments act as important biogeochemical hotspots for processing organic macromolecules. Interestingly, peptidase activity was negatively correlated to amounts of phyto-detritus inputs, whereas lipase activity was positively correlated [11]. This indicated that peptidases are excreted in times of nutrient limitation, whereas lipases are excreted when organic matter availability is high [11].

The microbial degradation of organic macromolecules in anoxic marine sediments is a complex inter-species process involving ‘primary degraders’ that break down larger macromolecules into oligomers and monomers for fermentation to alcohols, lactate and/or short-chain volatile fatty acids (VFAs), which are then mineralized to CH_4_, CO_2_ and/or H_2_ [21]. ‘Terminal oxidizers’ of fermentation products prevalent in marine sediments, such as sulfate-reducing microorganisms (SRM), have been relatively well studied [22–27]. On the other hand, the key microbial players responsible for the primary hydrolysis of different types of organic matter and macromolecules are poorly understood. Most studies are based on predictions from genomic analyses [28–31], whereas experimental evidence linking identities to functions are lacking.

In the present study, we investigated the identities, genomic features, and ecological interactions of microbial taxa that may play important roles in protein and lipid macromolecule degradation in subarctic marine sediments. We performed laboratory experiments whereby sulfidic arctic marine sediments were incubated in microcosms under cold (4 °C), anoxic conditions and supplemented with either ^13^C-labelled proteins or ^13^C-labelled lipids. The cold conditions represented large depth expanses of the oceans and seafloor that are permanently cold [32]. These microcosms were also incubated under conditions where sulfate reduction was specifically inhibited, in order to elucidate the roles of SRM. Catabolism and assimilation of ^13^C-labelled substrates by the sediment microorganisms was investigated by DNA-based stable isotope probing (DNA-SIP) and amplicon sequencing of 16S rRNA genes. The degradation of organic matter was monitored by measuring concentrations of VFAs and sulfate. Genome-resolved metagenomic analyses were used to predict secreted hydrolytic enzymes and reconstruct organic matter degradation pathways encoded by the taxa identified by DNA-SIP. This revealed (i) the identities and foraging strategies of several key protein- and lipid-consuming bacteria in marine sediments, (ii) that niche partitioning among *Psychromonas* subspecies was based on differential utilization of proteins and lipids, and (iii) and that SRM of the family *Desulfobacteraceae* mainly utilized the macromolecule degradation intermediates, i.e., volatile fatty acids.

## Materials and methods

### Sediment incubations

Sediment slurries were produced with sediment (0-30 cm below seafloor) from Greenland (Nuuk fjord ‘station 3’, water depth 498 m, August 2013, 64°26′45″N, 52°47′39″W) mixed 1:1 (v/v) with anoxic artificial seawater [33] containing approximately 28 mM sulfate. For each microcosm, 40 ml of this slurry was distributed into 250 ml serum vials under anoxic conditions in an anoxic glove-box (nitrogen atmosphere containing approximately 2% hydrogen and 10% CO_2_). Microcosms were sealed with thick rubber stoppers and crimped, and flushed with N_2_ after removing from the anoxic glove-box. All subsequent subsampling was also performed within the anoxic glove-box, with microcosms placed on ice-packs to minimize warming of incubations. Triplicate microcosms were either supplemented with a single dose of: i) 300 μg C g^−1 13^C-labelled Algal lipid mixture-99 atom% ^13^C (ISOTEC, Sigma-Aldrich); ii) 300 μg C g^−1 13^C-labelled Algal crude protein extract-98 atom% ^13^C (ISOTEC, Sigma-Aldrich), iii) 300 μg C g^−1 13^C-labelled Algal lipid mixture (as above) and the sulfate reduction inhibitor molybdate (28 mM; [34]); v) 300 μg C g^−1 13^C-labelled Algal protein mixture (as above) and molybdate (28 mM); or v) were left without supplementation (no-substrate controls). The microcosms were incubated at 4 °C and were sampled after 2, 5, 10, 17, 25 and 48 days of incubation. Sediment slurry samples were taken for analysis of volatile fatty acids and sulfate (1 ml), as well as for DNA-based analyses (0.5 ml). Samples were stored on pre-cooled ice packs in the anoxic glove box and were transferred immediately to the −80 °C freezer after sampling. Sampling, processing and molecular biological analyses of sediment samples from Svalbard, Norway, are presented as Supplementary information.

### Determination of sulfate and volatile fatty acid concentrations

Sulfate concentrations in interstitial water samples of microcosms were determined by capillary electrophoresis (P/ACETM MDQ molecular characterization system, Beckman Coulter) with the CEofixTM anions 5 kit (Analis). Standards were produced by dissolving known concentrations of Na_2_SO_4_ in artificial seawater. The concentrations of VFAs were determined as follows: samples were defrosted, vortexed and centrifuged at 10 000 rpm for 10 minutes. 100 ϋl of the supernatant was diluted 1:10 (v/v) in Milli-Q water. Syringe filters (Acrodisc IC grade filter, d=13 mm, Suprapor^®^ membrane with 0.2 μm pore size) were first rinsed with 10 ml Milli-Q water followed by 0.5 ml of diluted sample to minimize further dilution with the rinsing water. Another 0.5 ml of sample were then filtered and measured by 2-dimensional ion chromatography-mass spectrometry (IC-IC-MS; Dionex ICS-3000 coupled to an MSQ Plus™, both Thermo Scientific), equipped with an Ion Pack™ AS 24 as the first column to separate bulk VFAs from chloride, and an Ion Pack™ AS 11 HC as the second column to separate individual VFAs [35].

### DNA extraction

For DNA-SIP gradients and PCR-based amplicon sequencing, DNA was extracted using a combination of bead beating, cetyltrimethylammonium bromide-containing buffer and phenol-chloroform extractions. Sediment slurry (0.5 ml) was added to Lysing Matrix E tubes (MP Biomedicals) and were suspended in 0.675 ml cetyltrimethylammonium bromide containing extraction buffer, as described previously [36]. The tubes were placed on a rotary shaker (200 rpm) and incubated at 37 °C for 30 minutes. The samples were supplemented with 75 μL of sodium dodecyl sulfate (20% w/v) and were then incubated for 1 hour at 65 °C (tubes were inverted every 20 minutes). Samples were subjected to two rounds of bead beating with a speed setting 6 for 30 s using a FastPrep^®^-24 bead beater (MP Biomedicals). In between the beat beating steps, samples were cooled on ice. Debris was pelleted by centrifugation at 6000 g for 10 minutes at 25 °C and supernatant was collected. An equal volume of phenol:chloroform:isoamyl (25:24:1) alcohol (ROTH) was added to the supernatant and tubes were repeatedly inverted, followed by centrifugation at 16 000 g for 10 min at 25 °C. The aqueous phase (upper layer) was transferred into a clean 1.5 ml tube, supplemented with 0.6 volume of 2-propanol and incubated for 1 hour at 4 °C to precipitate DNA. The precipitate was then pelleted by centrifugation at 16 000 g for 30 min at 4 °C. DNA pellets were washed with 250 μL of 70% (v/v) ethanol, air-dried and resuspended in 50 μL Tris buffer [10 mM Tris-HCl (pH 8.0)]. For metagenome sequencing, DNA was extracted from day 0 and from individual sediment microcosm samples at days 5, 17 and 25 (for both protein and lipid amended treatments) using the Ultra Pure Power Soil Kit (MoBio) and following manufacturers’ protocol.

### DNA-SIP gradients

CsCl gradients with 5 μg DNA were prepared in a temperature controlled room at 23 °C according to a previously published protocol [37]. The gradient mixtures were added to ultracentrifuge tubes (Beckman Coulter) and were centrifuged in an Optima L-100XP ultracentrifuge (Beckman Coulter) using the VTi 90 rotor for >48 hours at 146286 rcf (44100 rpm) at 20 °C. After centrifugation, gradients were fractionated into 250 μl fractions by puncturing the bottom of the tube with a sterile needle and adding water to the gradient tube at the top using a sterile needle and a syringe pump (World Precision Instruments). The density of collected fractions was determined by using a digital refractometer (AR 200, Reichert Analytical Instrument) at 23 °C. The DNA-SIP fractions were considered heavy at densities >1.726 g ml^−1^ and light at densities <1.720 g ml^−1^ (Supplementary Figure S1). Afterwards, DNA was precipitated with 500mμL sterile PEG 6000 (30% polyethylene glycol 6000 and 1.6 M NaCl) and 1 μL of glycogen (5 μg ml^−1^) and subsequently purified as described previously [37]. Bacterial DNA in SIP fractions was quantified by quantitative PCR (qPCR) using the primers 341F 5′-CCT ACG GGA GGC AGC AG-3′ and 534R 5′-ATT ACG GCG GCT GCT GGC A-3′. The 20 μL qPCR mix contained 1× IQ™ SYBR Green Supermix (BIO-RAD), 0.25 μM of each primer and 1 μL of 1:10 diluted DNA from individual gradient fractions. The program used for thermal cycling on the iCycler thermal cycler (Bio-Rad) consisted of: 3 min at 95 °C, followed by 39 cycles of 95 °C for 15 s, 60 °C for 30 s and 72 °C for 39 s, and was followed by a melting curve from 60 °C to 95 °C by increments of 0.5 °C every 5 s.

### Amplicon sequencing and analysis

Barcoded 16S rRNA gene amplicons for Illumina MiSeq sequencing were produced using a previously established two-step PCR approach [38, 39]. Raw reads were processed as described previously [38, 39]. The identity threshold used for OTU clustering was 97% identity. Sub-OTU diversity at single-nucleotide resolution in the amplicon data was investigated using cluster-free filtering [40]. Statistical significance of differences in OTU relative abundances between treatments was determined using DESeq2 (version 1.10.1) [41] in the R software environment (R core team, 2019). Additionally, an OTU was considered to be ^13^C-enriched when it was significantly more abundant in heavy SIP fractions compared to light fractions obtained from the same incubation [42], which was also determined using DESeq2. The sequence abundance of each OTU in the ‘heavy’ part (>1.726 g ml^−1^) of the DNA-SIP gradient (numerator) were compared to sequence abundance in the ‘light’ part (<1.720 g ml^−1^) of the DNA-SIP gradient (denominator). Specific samples used for comparisons are indicated in Supplementary Figure S1. Only OTUs with more than 5 reads in 5 out of all 32 samples from gradient fractions were considered during DESeq2 analyses. Results were extracted with the command:

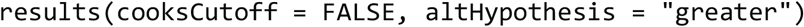

and were considered statistically significant if: (i) the false-discovery-rate (FDR)-adjusted p-value was below 0.1, and (ii) the respective OTU was not significantly enriched in the heavy fractions of the DNA-SIP gradient from the no-substrate incubations. OTU counts were ‘rlog’ transformed with DESeq2 and heatmaps of enriched OTUs were created with R software package pheatmap (version 1.0.8).

The presence of 16S rRNA gene sequences related (>97% identity) to specific organisms of interest in publicly available amplicon-derived datasets were determined using the IMNGS server [43], with 100 bp overlap as minimum.

### Metagenome sequencing and differential coverage binning

Metagenome libraries were produced with the Nextera XT DNA Library Preparation Kit (Illumina) and sequencing was performed at the Vienna Biocenter Core Facilities Next Generation Sequencing facility (Vienna, Austria) on an Illumina Hiseq 2500 using HiSeq V4 chemistry with the 125 bp paired-end mode. Reads were end trimmed at the first base with a q-score below 10 and reads with less than 50 bp were removed. For assembly, sequence coverage was normalized across samples using bbnorm with an average read depth of 100 and a minimum read depth of 3 (BBmap version 33.57 http://sourceforge.net/projects/bbmap/). Normalized read files were assembled with IDBA-UD [44] and SPAdes [45] using default parameters. Reads of each sample (non-normalized) were mapped to each assembly using BWA [46], and coverage information was obtained using SAMtools [47]. Differential coverage binning was performed with MetaBAT [48], MaxBin [49], and CONCOCT [50]. Following the binning programs default parameters, only contigs with a minimum length of 2500 and 1000 bp were used for binning with MetaBAT and MaxBin/CONCOCT, respectively. The resulting metagenome-assembled genomes (MAGs) generated from different assemblies and binning programs were aggregated with DASTool [51], and de-replicated with dRep using default parameters [52]. MAGs that could be linked to ^13^C-incorporating OTUs during protein and lipid hydrolysis by 16S rRNA gene sequence identity were further analysed. Selected MAGs were evaluated for completion and contamination using CheckM [53], and classified using the Genome Taxonomy Database (GTDB) release 86 [54] and GTDB-Tk version 0.1.3 [55]. A genome-based phylogenetic analysis of MAGs, the genome of the type strain *Psychrilyobacter atlanticus* HAW-EB21 and closely related reference genomes, was constructed from the concatenated marker protein alignment produced by CheckM [53]. The phylogenetic tree was built from this concatenated marker alignment with IQ-TREE using automatic substitution model selection (LG+F+I+G4) [56] and ultrafast bootstrap approximation with 1000 replicates [57]. The tree was visualized with iTOL [58]. Average nucleotide identities (ANI) and average amino acid identities (AAI) of selected MAGs and closely related reference genomes were determined using all-vs-all FastANI 1.1 with default parameters [59] and CompareM 0.0.23 (aai_wf) with default parameters (https://github.com/dparks1134/CompareM), respectively.

### Genome annotation

MAGs were annotated with the MicroScope annotation platform [60] and using the RAST server [61]. The annotations of proteins were confirmed by DIAMOND searches [62] against the NCBI-nr database (e-value 10^−5^), hidden Markov model-based searches using InterProScan [63] with the databases Pfam-A [64] and TIGRFAM [65], and online BLASTP searches against the UniProt database [66]. Possible hydrolytic enzymes were examined for signal peptide sequences that facilitate secretion from the cytoplasm and transmembrane helices using the Phobius online server [67] and using PSORTb version 3.0 [68]. Furthermore, peptidases, lipases/esterases and glycoside hydrolases were additionally compared to the databases MEROPS [69], ESTHER [70] and CAZY [71], respectively, using DIAMOND searches (e-value >10^−5^, identity ≥30%). Further, searches were made against the NCBI-nr database using BLASTP [72] for certain proteins of interest. The selection of potentially catabolic peptidases encoded by genomes and MAGs was based on two recent studies about extracellular peptidases in marine sediments [29, 73]. In addition, non-peptidase homologs and peptidases with regulatory functions, e.g., peptidases involved in membrane protein remodelling, as indicated by MEROPS database descriptions, were not considered (Supplementary Table S1). Potentially catabolic lipases/esterases were selected based on the ESTHER database descriptions (Supplementary Table S1).

### Phylogenetic analyses of 16S rRNA gene sequences

16S rRNA gene sequences were aligned and classified with the SINA aligner [74], using the SILVA database release 128 [75]. Full-length sequences of close relatives of 16S rRNA OTUs were extracted from the SILVA database and combined with the aligned 16S rRNA genes sequences that were obtained from MAGs. A reference tree was then constructed using FastTree [76]. Subsequently, 16S rRNA OTUS and partial 16S rRNA genes from MAGs were added to the reference tree using the EPA algorithm [77] in RAxML [78]. Trees were visualized in iTOL [58].

### Statistical analyses

Statistics for comparisons of VFAs and sulfate in treatment series versus controls were determined with Student’s T tests using the function t.test() in the R software environment (R core team, 2014).

### Data availability

All sequence datasets and metagenome-assembled genomes from the DNA-SIP experiments are available under the NCBI-Genbank Bioproject PRJNA609450. DNA-SIP amplicon sequencing datasets are available in the NCBI-Genbank Sequence Read Archive under BioSample accession SAMN14253696 and SRA accessions SRR11221408-SRR11221561. Metagenomic sequence reads are available in the NCBI-Genbank Sequence Read Archive under BioSample accession numbers SAMN14421543-SAMN14421548. Metagenome-assembled genomes are available in the NCBI-Genbank under BioSample accession numbers SAMN14421524-SAMN14421532. Annotated MAGs and genomes from the MicroScope annotation platform for key protein- and lipid-degrading organisms, and *Psychrilyobacter atlanticus* DSM 19335, are publicly available in the MaGe-Microscope server (https://mage.genoscope.cns.fr/). Amplicon sequencing datasets from Svalbard sediments are available under NCBI-Genbank Bioproject PRJNA623111 and the BioSample accession numbers SAMN14538997-SAMN14539076.

## Results

### Sulfate removal and volatile fatty acid turnover during protein and lipid degradation

Sulfate was largely turned-over after 48 days in all incubation treatments (down to 0.6-2.4 mM), except in incubations where sulfate reduction was inhibited by molybdate, where it remained between 18.1-24.2 mM (Figure 1A). Sulfate turnover was fastest between days 5 and 25, and was stimulated by the additions of proteins or lipids. Of all measured VFAs, acetate was the most prominent, reaching concentrations of over 900 μM in protein amended microcosms on day 17. Formation of the next most abundant VFA, formate, peaked at around 50 μM in lipid/molybdate-amended microcosms. Supplementation of proteins to the incubations resulted in significantly higher concentrations of acetate from days 2-5 compared to no-substrate controls (Supplementary Table S2). While the stimulation of acetate production from lipid additions was noticeably more than in no-substrate controls, the differences were not statistically significant (Supplementary Table S2). Significantly more propionate, butyrate and isobutyrate were produced in protein- and lipid-amended microcosms, as compared to the no-substrate controls (mainly between 2-5 days) (Figure 1B, Supplementary Table S2). Molybdate-inhibited incubations supplemented with proteins or lipids showed increased accumulation of formate, butyrate, and isobutyrate, but decreased production of acetate and propionate.

**Figure 1.**
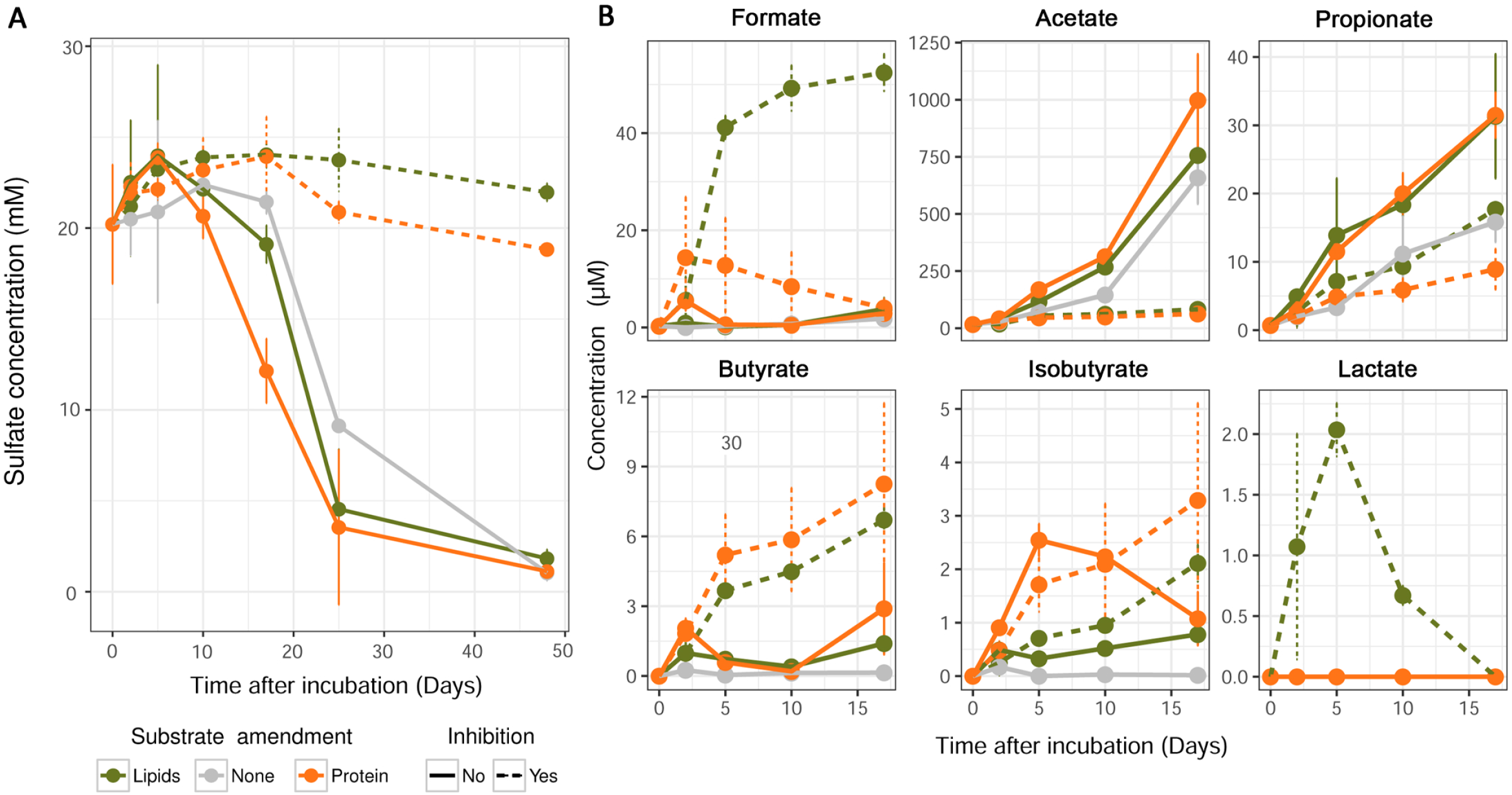
Concentration profiles of sulfate and fermentation products in subarctic marine sediment microcosms. Concentrations of sulfate (A) and volatile fatty acids (B) were determined in anoxic incubations with ^13^C-protein or ^13^C-lipids, and with or without the sulfate reduction inhibitor molybdate. Control microcosms were incubated without organic substrates and without molybdate. Error bars indicate standard deviation.

### ^13^C-labeling of OTUs and sub-OTUs during protein and lipid degradation

To evaluate incorporation of ^13^C from the added ^13^C-labelled proteins or lipids into the DNA of microorganisms, bacterial 16S rRNA genes were quantified by qPCR across individual SIP fractions (Supplementary Figure S1). Higher relative copy numbers were detected in the ‘heavy’ fractions of the gradients (i.e., densities >1.726 g ml^−1^) from both ^13^C-protein and ^13^C-lipid incubations relative to corresponding no-substrate controls at days 5 and 10. This indicated ^13^C-uptake and incorporation into DNA at these early time points. At days 17 and 25, only very small or no differences in relative 16S rRNA gene copy numbers between gradients from the ^13^C-substrate incubations and the no-substrate control incubations were observed.

Microbial community analysis of SIP fractions by 16S rRNA gene amplicon sequencing identified five OTUs (Clostridia-JTB215 OTU 1, *Psychromonas* OTU 4, *Psychrilyobacter* OTU 5, *Fusibacter* OTU 38 and *Photobacterium* OTU 54) that were significantly enriched in heavy SIP fractions from the ^13^C-protein incubations at day 5 (Figure 2A; Supplementary Table S3). At day 10, another 19 OTUs were significantly enriched in heavy SIP fractions from incubations with ^13^C-protein amendments (Figure 2A). These were affiliated with the classes *Deltaproteobacteria* (OTUs 2, 19, 36, 44, 67, 183, 232, 892, 4719), *Gammaproteobacteria* (OTUs 80, 128, 205, 285) and the phyla *Bacteroidetes* (OTUs 124 and 184), *Firmicutes* (OTUs 202 and 4050) and *Marinimicrobia* (OTU 123), or were unclassified (OTU 312) (Supplementary Figure S2).

**Figure 2.**
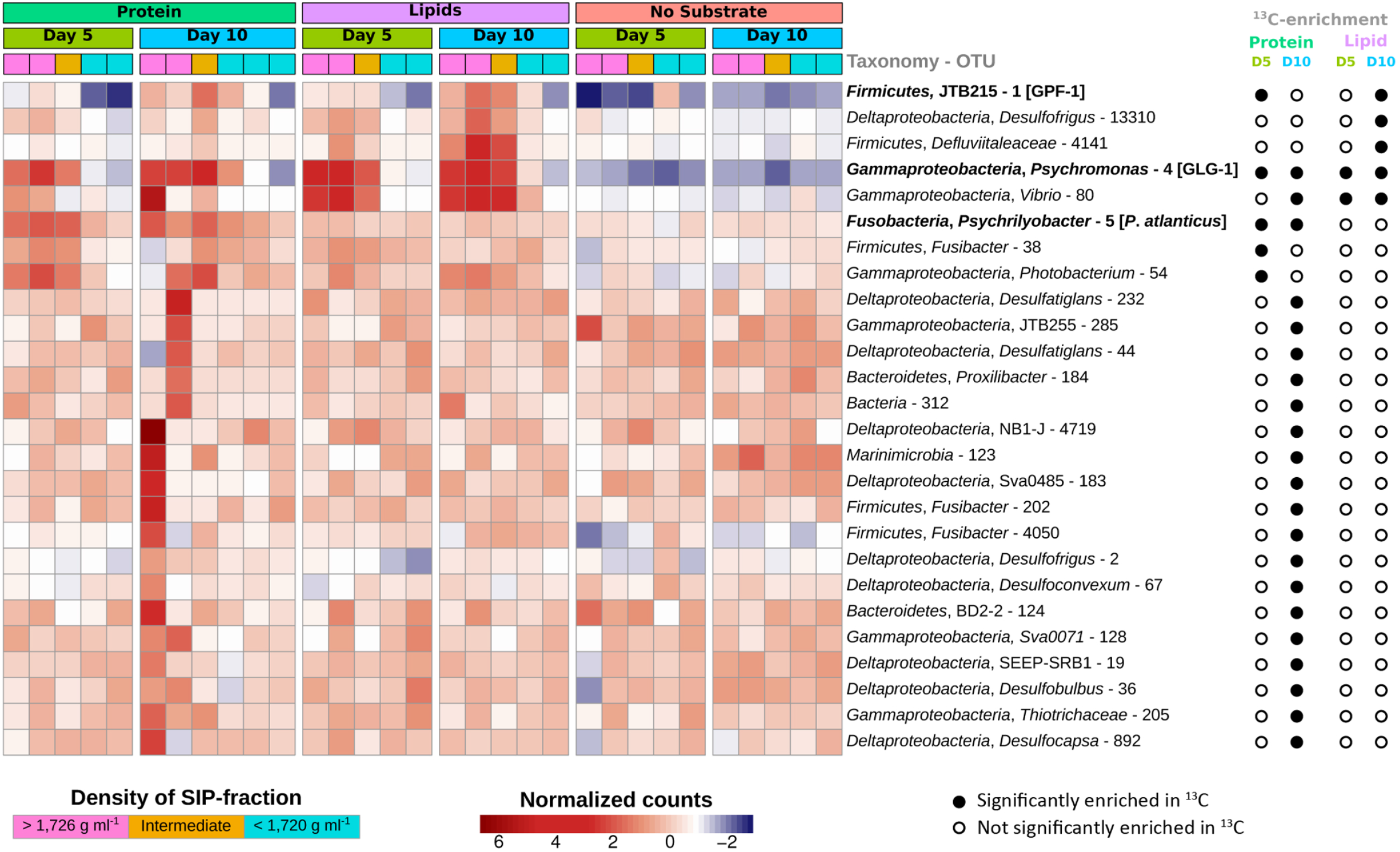
16S rRNA gene OTUs enriched in heavy ^13^C-DNA-SIP gradient fractions. Heatmaps showing the relative abundance of 16S rRNA gene OTUs in DNA-SIP gradient fractions from anoxic sediment incubations amended with ^13^C-proteins or ^13^C-lipids and from unamended control incubations. Significant (FDR-adjusted p-value was <0.1) ^13^C-enrichment was determined by differential abundance analysis between high density (^13^C-enriched, >1.726 g ml^−1^, pink blocks) and low density (^13^C-free, <1.720 g ml^−1^, turquoise blocks) DNA, and is indicated by filled circles. 16S rRNA genes that were matched to 16S rRNA gene sequences of genomes/MAGs are indicated in bold.

In SIP gradients from ^13^C-lipid amended incubations at day 5, significant ^13^C-enrichment was revealed for only two gammaproteobacterial OTUs, i.e., *Psychromonas* OTU 4 and *Vibrio* OTU 80 (Figure 2A). An analysis of the microdiversity within *Psychromonas* OTU 4 identified four *Psychromonas* sub-OTUs (sub-OTUs 4, 192, 9 and 42) that responded differently in incubations with protein or lipid amendments. The sub-OTUs 4 and 192 were significantly enriched in heavy fractions of ^13^C-lipid incubations (Figure 3A) and were more closely related to each other than to sub-OTUs 9 and 43 (Figure 3B). Furthermore, the relative abundance of sub-OTUs 4 and 192 increased in the microcosms from ^13^C-lipid incubations compared to no-substrate control microcosms (Figure 3C). In contrast, the sub-OTUs 9 and 43 were significantly enriched in heavy SIP fractions from ^13^C-protein incubations, and sub-OTU 9 increased in relative abundance in the microcosms from ^13^C-protein incubations compared to no-substrate control microcosms (Figure 3C). No sub-OTU microdiversity was detected in any other OTU that was significantly enriched in ^13^C. At day 10, we identified three additional OTUs enriched in the heavy SIP fractions from ^13^C-lipid incubations that were affiliated with the phylum *Firmicutes* (OTUs 1 and 4141) and the class *Deltaproteobacteria* (OTU 13310) (Figure 2A).

**Figure 3.**
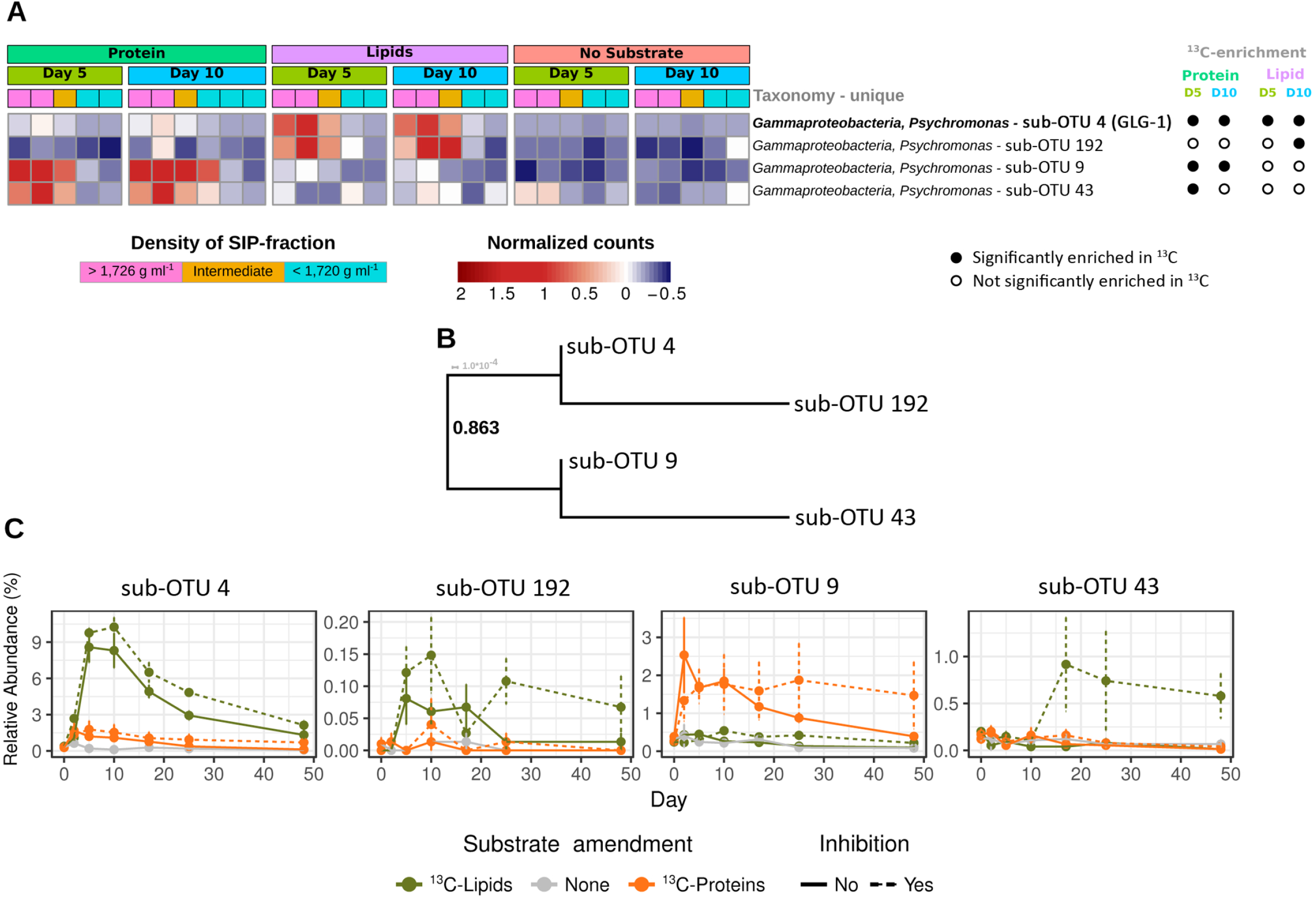
Differential ^13^C-labeling and response of *Psychromonas* sub-OTUs in protein- and lipid-amended sediment microcosms. (A) Heatmap showing the relative abundance of four sub-OTUs of *Psychromonas* 16S rRNA OTU 4 in DNA-SIP gradients. Significant ^13^C-enrichment was determined by differential abundance analysis between high density (^13^C-enriched) and low density (^12^C-enriched) DNA-SIP fractions. Sub-OTU 4, which was identical to the 16S rRNA gene sequence of *Psychromonas* MAG GLG-1, is indicated in bold. (B) Phylogeny of *Psychromonas* sub-OTUs. The sequences were aligned with MAFFT (Katoh et al., 2002) and the tree was calculated with FastTree (Price et al., 2010). (C) Changes in relative abundance of *Psychromonas* sub-OTUs in protein-amended, lipid-amended, and unamended sediment microcosms with or without the sulfate reduction inhibitor molybdate.

We also compared the relative abundances of 16S rRNA OTUs among the protein- or lipid-amended microcosms versus no-substrate control microcosms over time. Here, we restricted the analysis to OTUs (*n*=26) that were enriched in ^13^C from the protein- and lipid-amended treatments (described above). Between days 5 and 10, ten OTUs (1, 2, 4, 5, 19, 54, 80, 285, 4141 and 13310) significantly increased in relative abundances in substrate-amended incubations as compared to the no-substrate controls (Supplementary Figure S3A and Supplementary Table S2). Most notably, several OTUs showed relatively large differences in relative abundances between amended and no-substrate controls, and consequently had high relative abundances among the overall communities. For instance, *Psychrilyobacter* OTU 5 reached relative abundances of around 9-10% after 2-5 days in protein-amended microcosms, while staying around 3-4% in no-substrate controls. *Psychromonas* OTU 4 reached relative abundances of around 15-16% between 5-10 days after lipid amendments, while also reaching around 5-7% in the protein-amended microcosms from days 2-10. In contrast, *Psychromonas* OTU 4 stayed below 2.5% in the no-substrate controls over-time. The relatively fast (<10 days) and clear increases in relative abundances of these OTUs due to substrate additions therefore provided evidence for their direct involvement in substrate utilization for growth. On the contrary, the Clostridia-JTB215 OTU 1 also showed large increases in relative abundances from both protein- and lipid-amendments versus no-substrate controls, however, the response was relatively delayed, i.e., such increases developed mainly after 10 days.

Relative abundances of OTUs 1, 2, 67, 4050 and 4141 were significantly lower in microcosms with the sulfate-reduction-inhibitor molybdate than in microcosms without molybdate. This suggests that these OTUs are SRM or dependent on SRM activity.

### Genomic evidence for extracellular hydrolysis of proteins and lipids

Samples from day 0, and from days 5, 17 and 25 from the lipid- and protein-amended microcosms were selected for metagenome sequencing. The aim was to recover genomes of organisms that were ^13^C-labelled from protein or lipid treatments, and to specifically analyse their catabolic potentials for these macromolecules. Overall, nine metagenome-assembled genomes (MAGs) were recovered with >80% completeness and <5% contamination. Two had 16S rRNA sequences identical to bacterial OTUs that incorporated ^13^C from protein or lipid hydrolysis, i.e., *Psychromonas* MAG GLG-1 and *Clostridia* MAG GPF-1 (Supplementary Figure S2). All MAGs that were affiliated with the genus *Psychrilyobacter* were very incomplete (<20%) (data not shown). Therefore, we analysed the publicly available genome of the type strain *Psychrilyobacter atlanticus* HAW-EB21. This strain was isolated from marine sediments [79], and had a 16S rRNA sequence that was identical to *Psychrilyobacter* OTU 5 that was labelled from protein amendments (Supplementary Figure S2). A *Desulfoluna* MAG GLD-1 was also analysed in detail because it encoded potential to degrade VFAs and/or lipids, although it could not be confidently linked to any 16S rRNA OTU sequences and it is therefore presented in the Supplementary results.

*Psychromonas* MAG GLG-1 was most closely related to *Psychromonas aquimarina* ATCC BAA-1526 (ANI 80.7% of 41.8% aligned) (Supplementary Table S4) and encoded a 16S rRNA sequence that was identical to *Psychromonas* sub-OTU 4 (Supplementary Figure S2). *Clostridia* MAG GPF-1 was only distantly related to any publicly available genome (Supplementary Figure S4), with an AAI of only 53.4% (of 37% aligned) (ANI was too low to calculate) to the most-related genome of *Caloranaerobacter azorensis* DSM 13643 (Supplementary Table S4). A short 16S rRNA fragment (92 bp) was retrieved from *Clostridia* MAG GPF-1 and was linked to the 16S rRNA OTU 1 by phylogenetic placement into a reference tree (Supplementary Figure S2). The genomes/MAGs of putative protein and/or lipid-degraders were subsequently analysed for encoded capabilities to degrade proteins (*P. atlanticus* corresponding to 16S rRNA OTU 5), lipids (*Psychromonas* MAG GLG-1 corresponding to sub-OTU 4) or both macromolecules (*Clostridia* MAG GPF-1 corresponding to 16S rRNA OTU 1).

The genome of *Psychrilyobacter atlanticus* HAW-EB21 encodes a variety of peptidases (*n*=24), which were likely involved in the breakdown of peptides for further catabolism of amino acids (Supplementary Table S5). Two of these peptidases (i.e., M3 and M24) encoded signal peptides for translocation across the cytoplasmic membrane (Figure 4). Overall, the genome of *P. atlanticus* encodes an array of predicted peptide (*n*=19) and amino acid transporters (*n*=38), especially when compared to the MAGs of the putative lipid degraders *Psychromonas* MAG GLG-1 (peptide transporter, *n*=6; amino acid transporter, *n*=14) (Supplementary Table S5). Furthermore, the genome of *P. atlanticus* encoded for proton-dependent peptide transportation (Supplementary Table S5).

**Figure 4.**
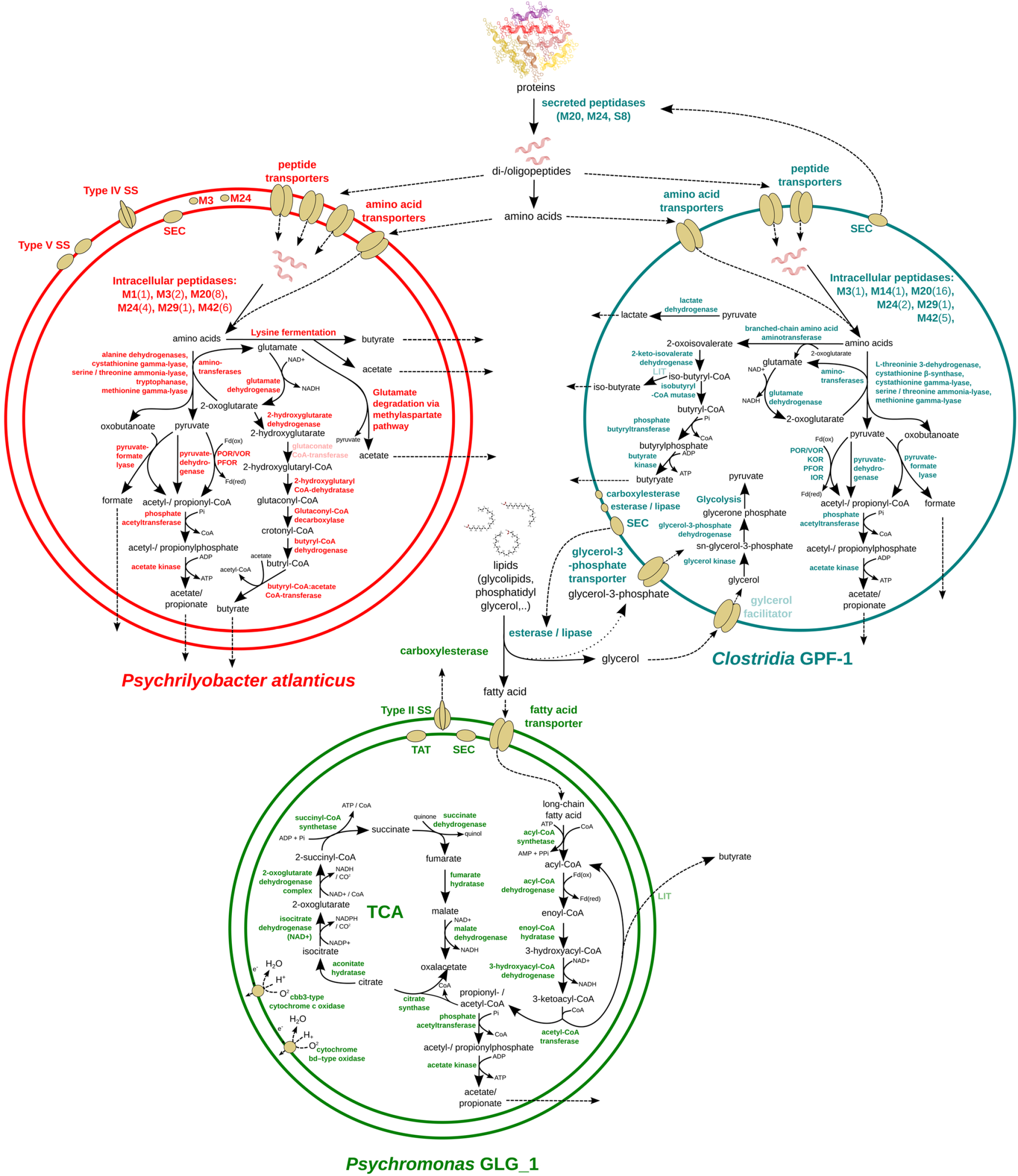
Genome-inferred metabolic models of protein- and lipid-degrading bacteria in arctic marine sediment. Schematic depiction of predicted metabolic properties of *Psychrilyobacter atlanticus*, *Clostridia* MAG GPF-1, and *Psychromonas* MAG GLG-1, which correspond to the 16S rRNA OTUs 5, 1, and 4, respectively. The models indicate macromolecule degradation processes and enzymes that may explain the observed biogeochemical patterns and the ^13^C-enrichment of corresponding 16S rRNA OTUs in the DNA-SIP incubations. Enzymes that are indicated in pale colour were not found in the respective genome/MAG. SS, Secretion system; TCA, tricarboxylic acid cycle; TAT, Twin-arginine translocation pathway; SEC, Sec secretion system; LIT, process has physiological evidence but the involved enzymes are not identified yet.

The lipid-metabolizing capability of *Psychromonas* MAG GLG-1 was supported by the presence of genes for an extracellular phospholipase D/alkaline phosphatase and a carboxylesterase with a signal peptide sequence, which corresponded to the ESTHER family Carboxylesterase type-B (Supplementary Table S5). The long chain fatty acids that resulted from extracellular degradation of acyl chain esters were likely taken-up by cells via two predicted long-chain fatty acid transporters (Supplementary Table S5). Predicted glycerol transporters were not encoded in the genome of *Psychromonas* MAG GLG-1. Because our study revealed *Psychromonas* spp. to be potentially important lipid and protein degraders, we also investigated available genomes and/or MAGs (*n*=16) from publicly available databases in order to examine the prevalence of the encoded capacity to utilize extracellular proteins or lipids via extracellular hydrolases (Supplementary Table S6). This identified predicted secreted phospholipases in most genomes (*n*=10), while predicted secreted proteases/peptidases were restricted to 4 genomes (Supplementary Figure S6).

*Clostridia* MAG GPF-1 encoded the largest number of peptidases (*n*=29) of all analysed MAGs/genomes (Supplementary Table S5). Three of these peptidases (M20, M24 and S8 peptidase families) have predicted secretion signal peptides and might be excreted to the extracellular space via the SEC pathway (Figure 4). Similar to *P. atlanticus*, *Clostridia* MAG GPF-1 encoded a broad array of predicted peptide (*n*=14) and amino acid (*n*=24) transporters. In accordance with the DNA-SIP results, *Clostridia* MAG GPF-1 also has the potential for catabolic breakdown of lipids. The MAG encoded one potentially secreted catabolic esterase/lipase that correspond to the ESTHER family Bacterial_EstLip_FamX, and two predicted membrane-bound esterases/lipases that correspond to the ESTHER families Bacterial_EstLip_Fam and Carboxylesterase, type B (Supplementary Table S5). The presence of genes for esterases/lipases in the genome of *Clostridia* MAG GPF-1 indicated the potential of this MAG to hydrolyse ester bonds between glycerol and fatty acids. Because genes encoding for known fatty acid transport and catabolism were absent from the genome of *Clostridia* MAG GPF-1, this bacterium might instead utilize glycerol. This was supported by an encoded glycerol-3-phosphate transporter and a putative glycerol facilitator protein, which were also encoded directly adjacent to a glycerol kinase, which catalyses the first step of the cytoplasmic catabolism of this substrate (Supplementary Table S5).

### Intracellular catabolism of protein and lipid degradation intermediates

The genome of *P*. *atlanticus* encoded 22 cytoplasmic peptidases with possible catabolic roles (Supplementary Table S5). In comparison, the genomes of the putative lipid degrader, i.e., *Psychromonas* MAG GLG-1 encoded only 7 intracellular peptidases. In this study, we focused on amino acid degradation processes that form glutamate and/or keto-acids for central carbon metabolism and that could therefore explain incorporation of ^13^C-carbon from the ^13^C-labelled proteins into DNA, as well as the development of VFAs in our incubations (Figure 4). The genome of *P*. *atlanticus* encoded several aminotransferases and other amino acid degrading enzymes (i.e., alanine dehydrogenases, cystathionine gamma-lyase, serine/threonine ammonia-lyase, tryptophanase and methionine gamma-lyase) (Supplementary Table S5). These can convert amino acids to the keto-acids pyruvate and oxobutanoate, which could then be fermented to formate, and either acetate or propionate, respectively (Figure 4).

*Psychromonas* MAG GLG-1 encoded all enzymes required for beta-oxidation of fatty acids, including a putative multienzyme complex that contains the two enzymes enoyl-CoA hydratase/isomerase and 3-hydroxyacyl-CoA dehydrogenase (Supplementary Table S5). Through beta-oxidation, even- and odd-chain fatty acids can be converted to propionyl- and acetyl-CoA, which can be further fermented to acetate and propionate, respectively (Figure 4). In comparison to *P. atlanticus* and *Clostridia* MAG GPF-1, which encoded strict fermentative metabolisms, *Psychromonas* MAG GLG-1 also encoded a complete oxidative TCA cycle, which enables a respiratory conversion of acetyl-CoA to CO_2_ (Figure 4). When oxygen is available, *Psychromonas* MAG GLG-1 might use one of the two encoded high affinity oxidases for respiration, i.e., cbb3-type cytochrome *c* oxidase and cytochrome *bd*-type oxidase (Supplementary Table S5). In oxygen-depleted sediments like in our microcosm incubations, *Psychromonas* GLG-1 might ferment, or may respire using nitrate (dissimilatory nitrate reduction to ammonium, DNRA) or fumarate as terminal electron acceptors (Supplementary Table S5). Genes necessary for the intracellular degradation of glycerol were absent from *Psychromonas* MAG GLG-1 (Supplementary Table S5). Furthermore, despite the presence of genes for several extracellular glycosyl hydrolases in the genome of *Psychromonas* MAG GLG-1 and a complete glycolysis pathway (Supplementary Table S5), enzymes for feeding galactose or glycerol (subcomponents of Spirulina-derived galactosyl diglycerides), into glycolysis, were not identified.

The *Clostridia* MAG GPF-1 encoded the largest set of intracellular peptidases (*n*=26) of all investigated genomes (Supplementary Table S5). The fermentation of amino acids by *Clostridia* GPF-1 to propionate, acetate or formate, i.e., via aminotransferases and other amino acid degrading enzymes (i.e., L-threonine 3-dehydrogenase, cystathionine β-synthase, cystathionine gamma-lyase, serine/threonine ammonia-lyase and methionine gamma-lyase) was mostly similar to the predicted routes used by *P. atlanticus*, although genes for lysine fermentation were not present and the glutamate degradation pathways were incomplete (Supplementary Table S5). In addition, the *Clostridia* MAG GPF-1 was the only MAG that encoded enzymes that initiate valine degradation, i.e., two branched-chain amino acid transferases and the gene 2-keto-isovalerate dehydrogenase that is part of the branched-chain α-keto acid dehydrogenase complex (Supplementary Table S5). Besides the potential for amino acid degradation, *Clostridia* MAG GPF-1 also encoded all genes necessary for the conversion of glycerol and glycerol-3-phosphate to pyruvate (Supplementary Table S5), which may help explain with the ^13^C enrichment of 16S rRNA OTU 1 in ^13^C-lipid amended incubations (Figure 2).

### Environmental distributions of key taxa

To explore fine-scale depth distributions of the main primary-hydrolysing *Psychromonas* and *Psychrilyobacter* species in sulfidic arctic marine sediments, we examined 16S rRNA-gene and -cDNA derived sequence datasets from sediments of Smeerenbergfjord, Svalbard (Supplementary Figure S6). This showed that *Psychromonas* sequences reached 1.7% and 1.9% relative abundances in 16S rRNA-gene and -cDNA libraries in 0-1 centimetres below seafloor (cmbsf) samples from Station GK, respectively. In a nearby core from Station J, they also reached 1.2% relative abundances in both 16S rRNA-gene and -cDNA libraries in 0–1 cmbsf samples. At both sites, relative abundances of *Psychromonas* 16S rRNA-genes and -transcripts were very low under 3 cmbsf (Supplementary Figure S6). Sequences from the *Psychrilyobacter* reached only 0.3% in 0–1 cmbsf from Station J samples and comprised maximally only 0.05% of the 16S rRNA transcripts in the same sample (results not shown). *Psychrilyobacter* sequences were under 0.03% in all other samples deeper than 1 cmbsf. Further, to gauge their prevalence in other marine sediments, we examined sequences that were similar (>97% identity) to the *Psychromonas* and *Psychrilyobacter* OTUs recovered in this study to sequences in publicly available 16S rRNA gene datasets. This showed that sequences related to these organisms were widely distributed in sediment sites around the worlds oceans (Supplementary Figure S7). For example, *Psychromonas* sequences reached a maximum of 4% relative abundance in seagrass-associated sediments (Nantucket Sound, USA).

## Discussion

Proteins and lipids comprise a significant fraction (up to 10% each) of organic matter in marine sediments [8, 9], and therefore represent major nutrient and energy sources for sediment microbiomes. Our experiment showed that supplemented proteins and lipids were actively utilized by microorganisms within 5 days of incubation. Both protein and lipid amendments induced rapid accumulation of several VFAs, demonstrating primary fermentation of both macromolecules. DNA-SIP further showed that ^13^C-carbon from protein and lipid catabolism was incorporated into the DNA of specific taxa within 5 days. This was also paralleled by sharp increases in the relative abundances by many of the same taxa in the corresponding microcosms. We therefore reasoned that taxa that were ^13^C-labelled and stimulated to higher relative abundances (>1%) within the first 5 days represented the main primary hydrolysers and fermenters of proteins or lipids. Thus, the following discussion especially focuses on these primary hydrolysing taxa.

Overall, more diverse OTUs incorporated ^13^C-carbon from proteins compared to lipids, especially after 10 days of incubation, possibly due to a combination of: i) higher proportions and bioavailability of proteins in sediments [8, 9], ii) greater nutritional value of proteins (e.g., nitrogen), and/or iii) a greater biodegradability of proteins as compared to lipids [80]. Sediment microorganisms are also known to efficiently salvage organic nitrogen sources such as amino acids in times of low nutrient supply [11]. Therefore, some microorganisms may have salvaged free peptides or amino acids released from the primary hydrolysers and became ^13^C-labelled.

The *Psychromonas* 16S rRNA gene OTU 4 was notable in this study because our DNA-SIP results showed that it was a prominent degrader of both lipids and proteins. Interestingly, we could distinguish four sub-OTUs that each had distinct preferences for either lipids or proteins. Thus, this demonstrated that closely related populations of *Psychromonas* in marine sediments have very different preferences for macromolecules that require completely different catabolic machinery. From the metagenomic analyses, the *Psychromonas* MAG GLG-1, which represented the lipid-degrading sub-OTU 4 population, indeed had gene content that supported its capacity to digest lipids and catabolize fatty acids. For instance, it encoded a secreted esterase/lipase to digest lipids from the extracellular environment, as well as long-chain fatty acid transporters and a beta-oxidation pathway to import and catabolize the fatty acids (Figure 4). Although we did not recover any *Psychromonas* MAGs representative of the protein-degrading populations, comparative genomic analyses of publicly available *Psychromonas* genomes and MAGs showed predicted secreted lipases and/or peptidases are common among this genus. Diverse members of this genus therefore have unique potentials to utilize lipids or proteins.

Our microbial community profiling showed that *Psychromonas* were relatively abundant and active (based on 16S rRNA expression) in shallow sediments (<3 cmbsf) from arctic Svalbard. *Psychromonas* were previously shown to be highly correlated with fluxes of fresh phyto-detritus to surface sediments of the arctic Laptev Sea [81]. Together with our experimental and genomic analyses, this suggests *Psychromonas* play an active role in the turn-over of ‘fresh’ and labile organic matter delivered to surface marine sediments from the water column. This is inline with previous experiments that showed *Psychromonas* were important players in the initial degradation of whole Spirulina (cyanobacterial) necromass in arctic marine sediment [23]. Our study therefore suggests that *Psychromonas* populations thrive in surface sediments by utilizing abundant macromolecules that represent fresh and labile organic matter, e.g., lipids and/or proteins.

We further showed that *Psychrilyobacter* OTU 5 was one of the most prominent protein and/or amino acid degraders in our experiment. Members of this genus were also shown to be important degraders of whole Spirulina necromass in marine sediment experiments [23, 33]. Here, we therefore deciphered their role in bulk organic matter degradation by showing that they are efficient degraders of detrital protein/peptides. Correspondingly, genomic analyses showed that the *Psychrilyobacter atlanticus* type strain HAW-EB21 [79], which had 100% 16S rRNA sequence identity to OTU 5, encoded various routes for the degradation of peptides and for the subsequent fermentation of amino acids (Figure 4) [79]. Their overall contributions to protein degradation in marine sediments may, however, be less than the *Psychromonas* because they were not abundant members of most sedimentary microbiomes analysed.

The Clostridia MAG GPF-1 (16S rRNA-OTU 1), had less than 54% AAI to any available genome, and therefore our data represents the first insights into the potential metabolism of this uncultured group. Overall, our analyses suggested these organisms are versatile nutrient utilizers. The Clostridia OTU 1 population was ^13^C-labelled in SIP analyses from both ^13^C-protein and -lipid incubations, and its relative abundances were also boosted by the additions of both substrates, although to a much lesser extent than the *Psychromonas* or *Psychrilyobacter*. Our genomic analyses indicated the potential to utilize peptides, amino acids and glycerol (Figure 4), thereby highlighting its versatility. Further, OTU 1 also increased in no-substrate controls over time, suggesting it either utilized natural organic matter in the microcosms, or fermentation products produced from the degradation of the natural organic matter by other members of the community.

In addition to the relatively abundant taxa described above that were ^13^C-enriched and that could be linked to genomes or MAGs, we also identified ^13^C-enrichment in several low abundance taxa (<1% relative abundances), i.e., members of the genera *Photobacterium* (OTU 54), *Vibrio* (OTU 80) and *Fusibacter* (OTU 38). These taxa may have therefore played a lesser role in the primary hydrolysis of the macromolecules and are therefore discussed briefly in the Supplementary Discussion.

After the main primary degradation processes and increased formation of fermentation products (i.e., VFAs) in the first 5 days, ongoing depletion of sulfate and inhibition of sulfate depletion by molybdate indicated SRM were active. Various taxa including *Deltaproteobacteria* related to known SRM became ^13^C-labelled at day 10, which suggested transfer of carbon from the added macromolecules to SRM. This was likely via oxidation of the VFAs by SRM, because deltaproteobacterial SRM are known to be important oxidizers of fermentation products in marine sediments [82–84]. The detection of more ^13^C-labelled putative SRM of the *Deltaproteobacteria* from protein incubations than from lipid incubations may have reflected the more diverse taxa involved in primary degradation of the proteins, e.g., through more diverse metabolic interactions. We also hypothesize the additional nitrogen supply from the proteins may have facilitated the growth of more diverse *Deltaproteobacteria*, since none of the VFAs appeared to be limiting from either protein or lipid treatments.

In conclusion, this study provided new insights into the identities and functions of various protein- and lipid-degrading microorganisms in cold marine sediments and indicates *Psychromonas* spp. are prominent players in the utilization of fresh detrital protein and/or lipid macromolecules. The activity of these primary hydrolysers also facilitates relatively rapid transfer of carbon and energy among trophic levels within the sediment microbial community.

## Acknowledgements

The authors would like to thank the crew of the R/V Sanna 2013 summer sampling campaign, which was funded by the Arctic Research Centre, Aarhus University, and Kasper Kjeldsen, Hans Røy and Marit-Solveig Seidenkranz for providing the sediment samples. We thank captain (Stig Henningsen) and first mate of MS Farm for their assistance during sampling of Svalbard sediments in 2017, as well as the Kings Bay Marine Laboratory and the AWIPEV Arctic Research Base in Ny-Ålesund for assistance and laboratory space. Furthermore the authors want to thank Carmen Czepe from the Vienna Biocenter Core Facilities Next Generation Sequencing facility for sequencing of metagenomic libraries on the Illumina Hiseq2500 instrument. This work was financially supported by the Austrian Science Fund (P29426-B29 to KW; P25111-B22 to AL).

The authors declare no conflict of interest.

