## Supplementary information text for "Anaerobic microbial degradation of protein and lipid macromolecules in subarctic marine sediment"

**Supplementary materials and methods**

***Microbial community analysis of Svalbard samples***

Sediment samples from Smeerenburgfjorden, Svalbard, were collected in June 2017. The samples were taken with a HAPS corer from the vessel ‘MS Farm’ from station GK (79°38.49N 11°20.96E), and station J (79°42.83N 11°05.10E). The HAPS cores were subsampled on deck with a plastic subcorer that had 5 cm width and pre-drilled ports. Subsamples (1 mL of sediment) were taken with sterile 3 ml syringe, added to 2 ml tubes and flash frozen in a dry-shipper pre-cooled by liquid nitrogen. Sample were stored at -80°C in the laboratory until further analyses.

Extractions of RNA/DNA were performed with the RNeasy PowerSoil Total RNA Kit (Qiagen) according to manufacturer’s protocol. From the eluted nucleic acids, DNA for PCR was used as eluted. Aliquots of RNA extracts were DNase treated with the TURBO DNA-free kit (Thermo Fisher) following the manufacturer’s protocol. RNA was reverse transcribed using the Revert Aid First Strand cDNA Synthesis Kit (Thermo Fisher) following the manufacturer’s protocol. A reverse transcription negative control was performed by combining the remaining supernatant of the samples and adding all reagents except the Revert Aid M-MuLV Reverse Transcriptase. The cDNA was checked by PCR with primers targeting bacterial 16S rRNA genes and gel electrophoresis. The reverse transcription-negative control was always negative.

For 16S rRNA gene amplicon sequencing, a two-step PCR and barcoding approach was used [[1]](https://paperpile.com/c/TbafTS/FK1b). The primers 515F (5′-GTGYCAGCMGCCGCGGTAA-3′) and 806R (5′-GGACTACNVGGGTWTCTAAT-3′) were used in the first step, and both primers included a linker sequence to facilitate barcoding in the second-step PCR. All first-step PCRs were performed in triplicates (and combined after PCR for barcoding) with a reaction volume of 12.5 µl. Each reaction (12.5µl) contained: 1X Dream Taq Buffer (including 2 mM MgCl_2_) (Thermo Fisher), 0.2 mM dNTP mix (Thermo Fisher), 0.2 µM of each forward and reverse primer, 0.08 mg ml^-1^ BSA (Thermo Fisher), 0.02 U Dream Taq Polymerase (Thermo Fisher), UV-treated deionised water (up to 12 µl) and 0.5 µl of template. PCR cycling was: 95°C for 3 min, followed by 30 cycles of 30 sec denaturation at 95°C, 30 sec annealing at 52°C and 50 sec elongation at 72°C, and a final 10 min at 72°C. Sequencing was performed by the Joint Microbiome Facility (Vienna, Austria) using an Illumina MiSeq with MiSeq Reagent Kit v3 chemistry with 300 bp paired-end read mode. Bioinformatic processing of 16S rRNA gene amplicon data from Svalbard sediments was performed by demultiplexing amplicon sequencing variants (ASVs) that were constructed using DADA2 [[2]](https://paperpile.com/c/TbafTS/o0ie) with the previously described workflow [[3]](https://paperpile.com/c/TbafTS/PUBY).

**Supplementary results**

The deltaproteobacterial *Desulfoluna* MAG GLD-1 encoded the capacity to utilize various VFAs and potentially lipids. In addition to several acyl-CoA synthetases, it encoded a butanoate CoA-transferase that enables conversion of butyrate to butyryl-CoA, which can then be funneled into beta-oxidation (Supplementary Figure 8). For lactate degradation, GLD-1 encoded several lactate dehydrogenases and a lactate utilization operon (Supplementary Table 5). GLD-1 also encoded a formate dehydrogenase enzyme complex (Supplementary Table 5) to oxidize formate to CO_2_ and H_2_ (Figure 5). Furthermore, the potential to utilize lipids and long-chain fatty acids was indicated by genes for an outer membrane phospholipase A, a long-chain fatty acid transporter, and multiple long-chain fatty acid CoA ligases (Supplementary Table 5) with high similarity to homologs from other known long-chain fatty acid- and lipid-degrading SRM. The *Desulfoluna* MAG GLD-1 encoded the complete pathway for dissimilatory sulfate reduction (Supplementary Figure 8).

**Supplementary discussion**

In addition to the relatively abundant taxa described above that were ^13^C-enriched and that could be linked to genomes or MAGs, we also found ^13^C-enrichment in several less-abundant taxa (<1% relative abundances in microcosms). Members of the *Fusibacter* (OTU 38) and *Photobacterium* (OTU 54) were determined to be ^13^C-labelled from protein incubations, while a *Vibrio* species (OTU 80) was determined to be ^13^C-labelled from lipid incubations. The *Photobacterium* (OTU 54) and *Vibrio* (OTU 80) also showed increased relative abundances in microcosms that were also specific to protein or lipid amended microcosms, respectively. This supported their roles in primary hydrolysis of these substrates, or had specific associations with primary hydrolyzers of either substrate. Members of these groups have been previously described to have relatively diverse heterotrophic metabolisms. For instance, close relatives of *Photobacterium* OTU 54 were previously shown to be both lipolytic [[4, 5]](https://paperpile.com/c/TbafTS/57po6+Iq2Lu) and proteolytic [[6, 7]](https://paperpile.com/c/TbafTS/JkGCn+XYt1K). The majority of OTU 80 related *Vibrio* *spp*. are able to degrade sugars and glycerol, and also include lipolytic isolates [[8–13]](https://paperpile.com/c/TbafTS/z8OHv+oJsKX+MrYF2+Ii432+Vm81r+xPnE5). These data therefore suggest that these taxa may have played roles in primary hydrolysis of the macromolecules. In comparison, the *Fusibacter* (OTU 38) gradually increased in relative abundances in all microcosms including no-substrate controls, thereby suggesting they may have rather utilized fermentation products or degradation products common to the different treatments, e.g., acetate.
