## Supplementary Figures 1-8 for "Anaerobic microbial degradation of protein and lipid macromolecules in subarctic marine sediment"

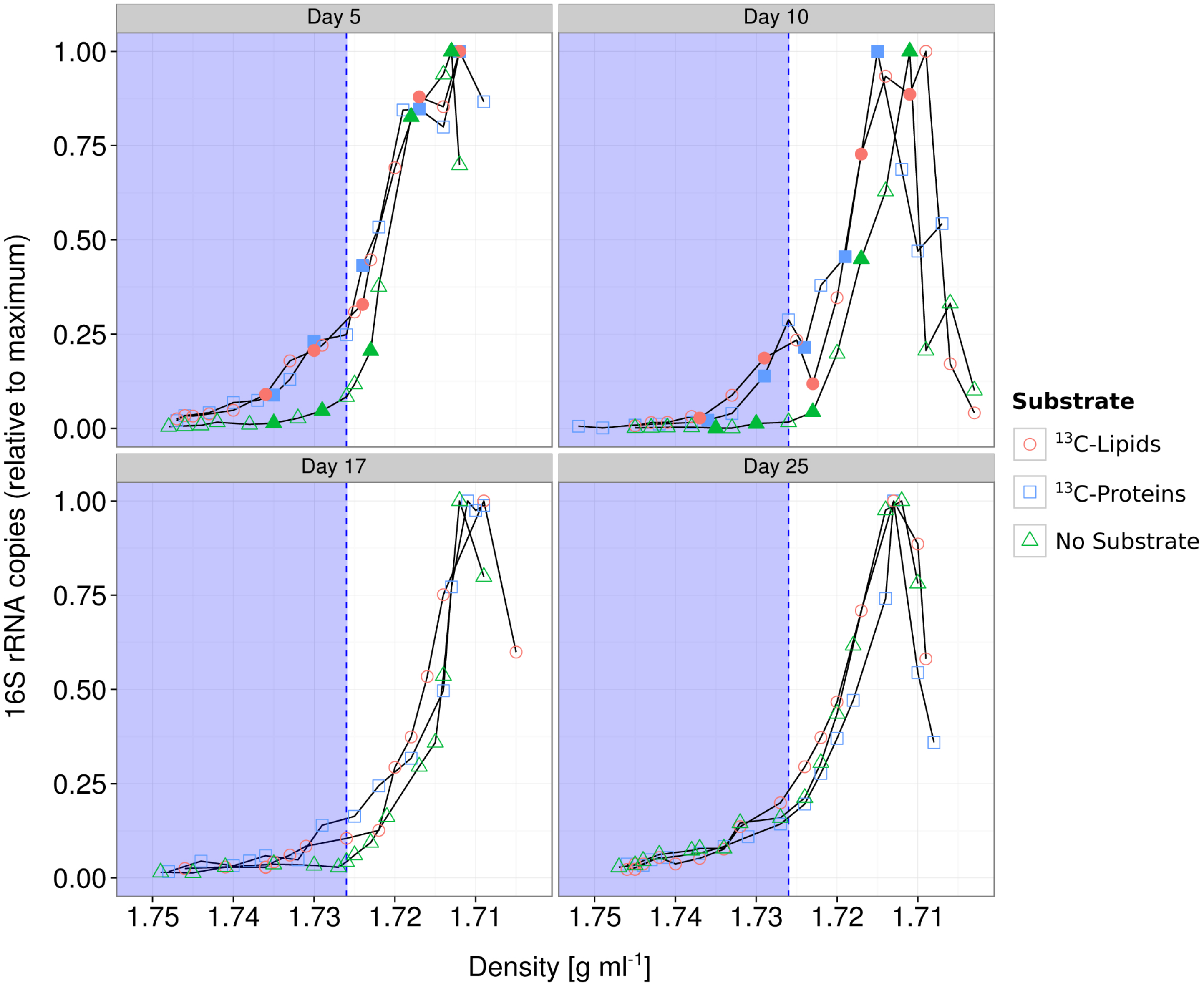


**Supplementary Figure S1. Proportions of 16S rRNA gene copies recovered from fractions of DNA-SIP gradients.** Color and shape of data points indicate the amended substrate. Filled symbols indicate that DNA from the respective fraction of the DNA-SIP gradient was used as template for amplicon sequencing. The blue shaded area indicates the density of the gradient above which $^{13}C$-labelled DNA is expected to accumulate.

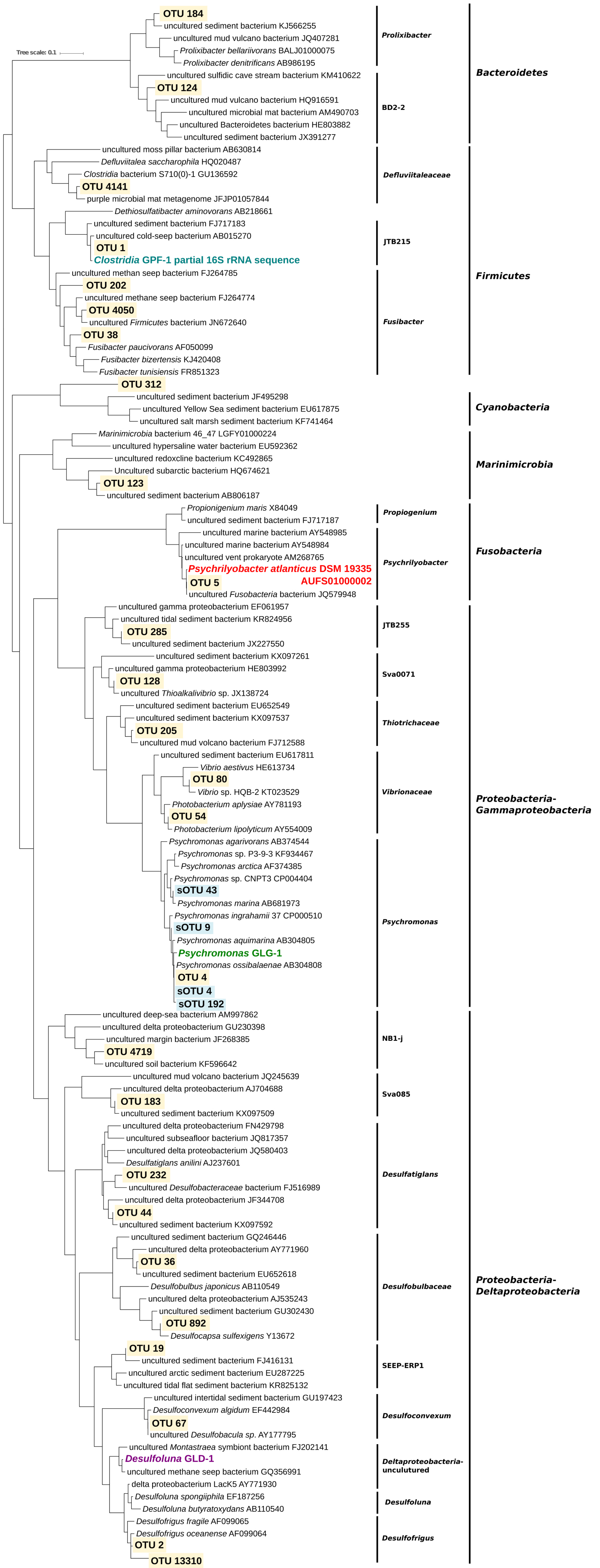


**Supplementary Figure S2. Phylogenetic affiliation of 16S rRNA OTU sequences.** Tree was constructed from close relative sequences of SIP-enriched OTUs in thw SILVA database v.128 (Quast *et al.*, 2013) and full-length 16S rRNA genes from MAGs with Fasttree (Price *et al.*, 2010). Full-length MAG derived 16S rRNA genes were aligned to the SILVA database v.128 (Quast *et al.*, 2013) with the SINA aligner (Pruesse *et al.*, 2012) prior to tree construction. 16S rRNA OTUs and partial 16S rRNA genes from MAGs were aligned to the SILVA database and then placed into the reference tree using the EPA algorithm (Berger *et al.*, 2011) in RAxML (Stamatakis, 2014). The tree was visualized  iTOL (Letunic and Bork, 2007).

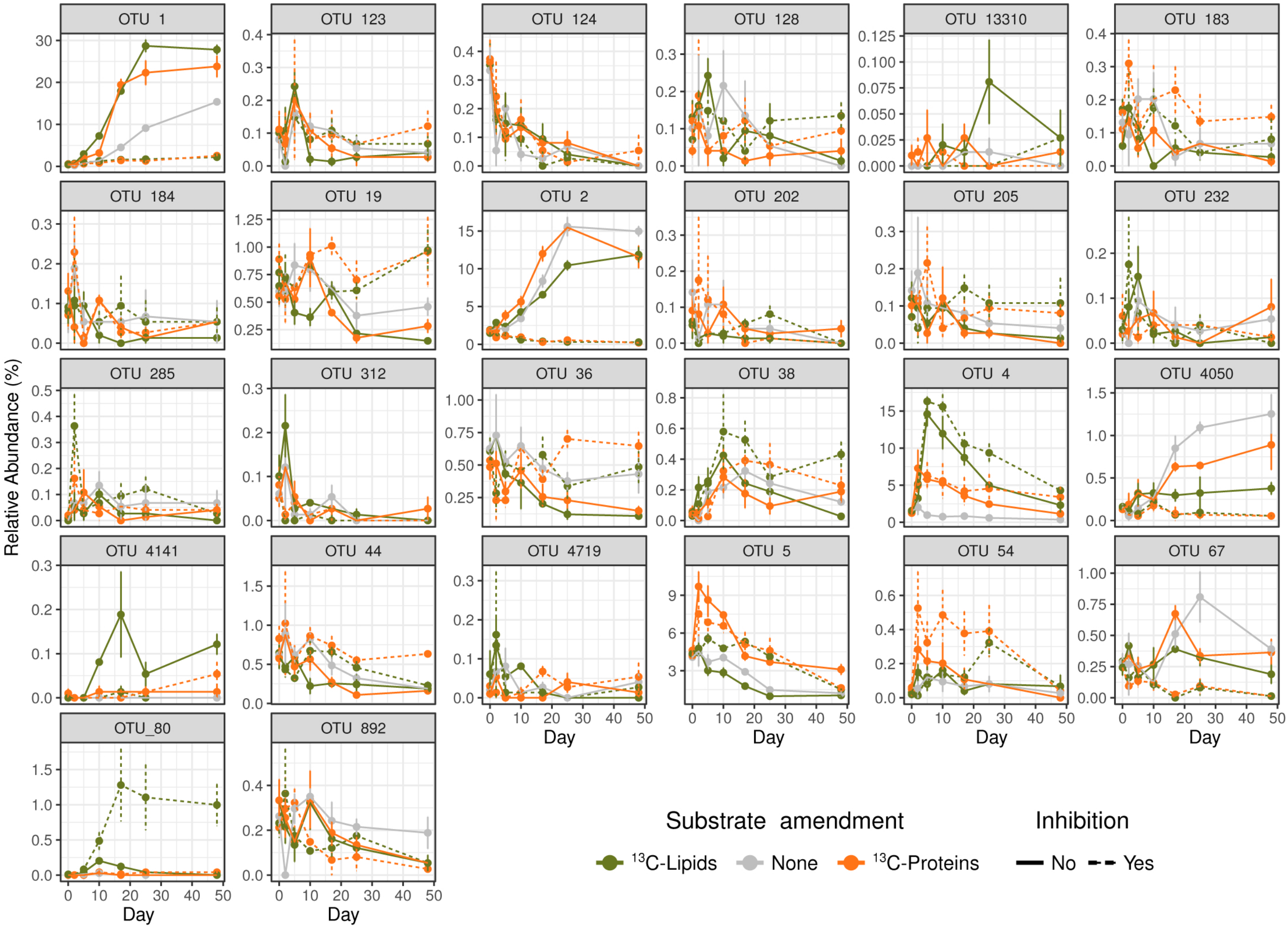


**Supplementary Figure S3**. Relative abundances of OTUs that incorporated $^{13}C$-carbon over-time in microcosms. Inhibition = microcosms where molybdate was added to inhibit sulfate reduction.

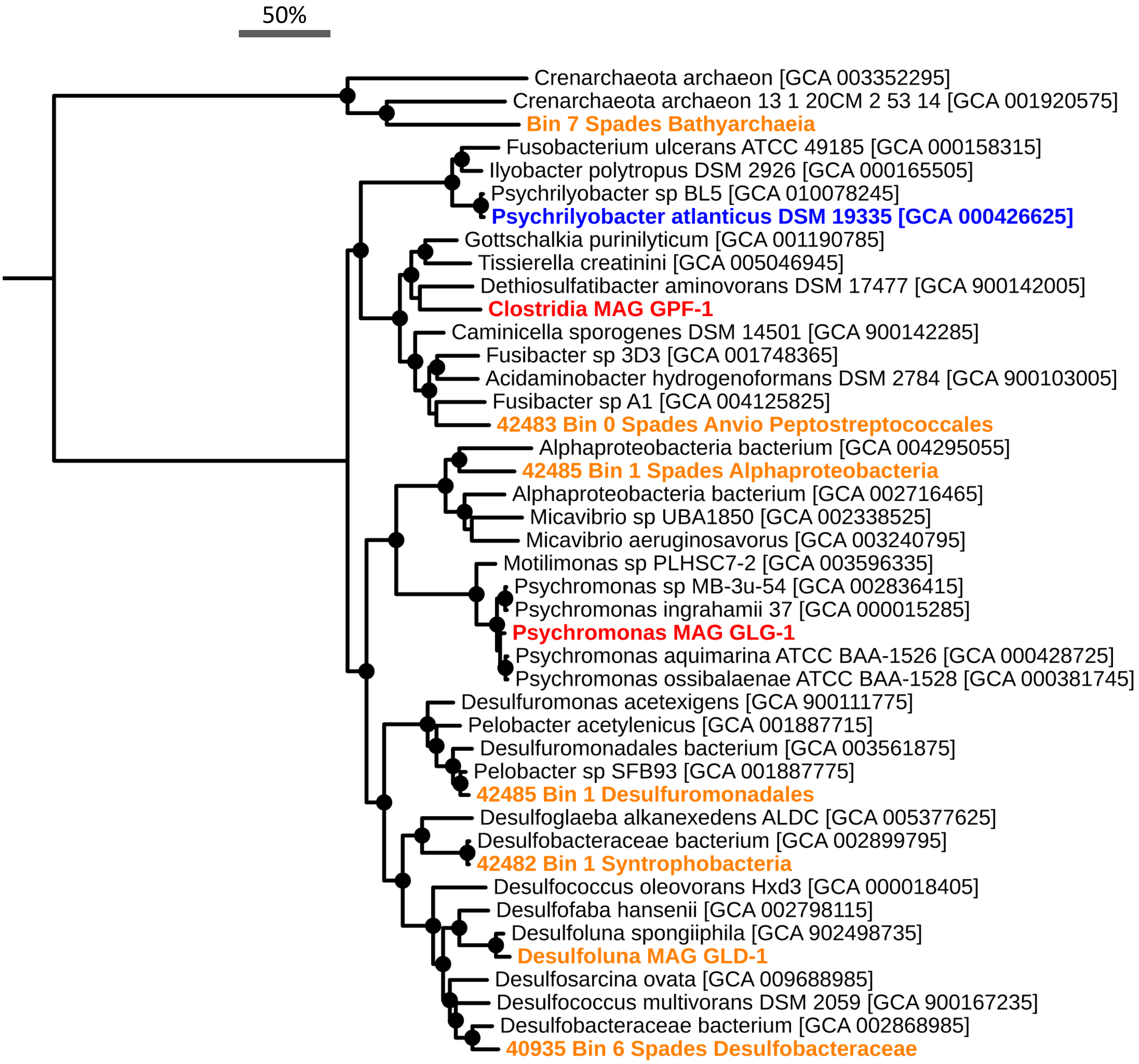


**Supplementary Figure S4. Phylogenetic analysis of concatenated single copy marker proteins.** The phylogeny (maximum likelihood) is based on concatenated protein sequences derived from single copy marker genes retrieved from CheckM analyses. Red leaves correspond to MAGs of organisms determined to be labelled from DNA-SIP. The blue leaf corresponds to the genome of reference sequence *Psychrilyobacter atlanticus* DSM 19335. Orange leaves correspond to other MAGs recovered in this study. Genbank Bioproject accesssion numbers for MAGs from this study are presented in Supplementary Table S4. Genbank assembly accession numbers for reference genomes are presented in parenthesis. Bootstrap values of >90% are indicated by filled black circles. The scale bar represent 50% sequence divergence.

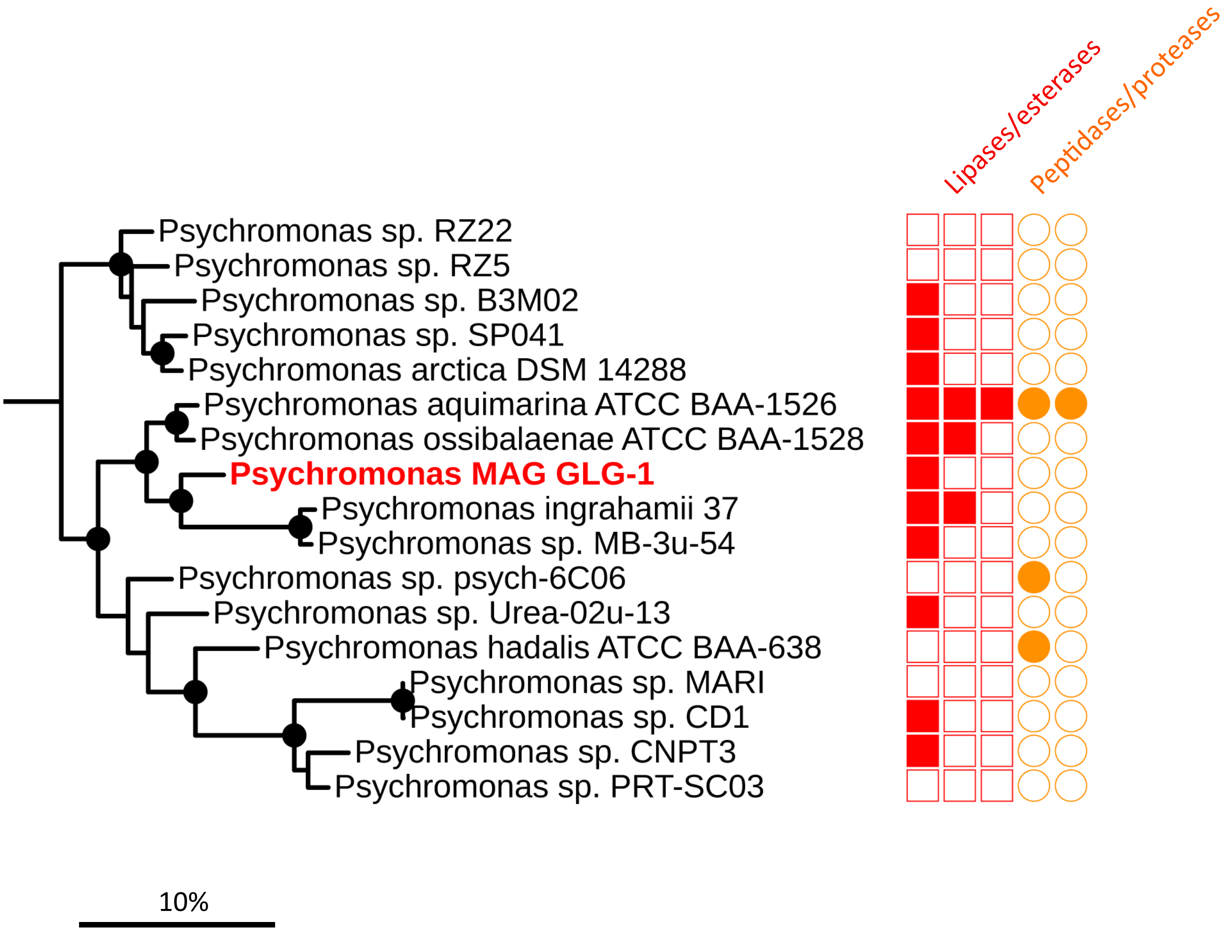


**Supplementary Figure S5**. Phylogenomic tree of available *Psychromonas* spp. genomes and MAGs, and *Psychromonas* MAG GLG-1 (bold red) recovered in this study. The phylogeny (maximum likelihood) is based on concatenated protein sequences derived from single copy marker genes retrieved from CheckM analyses. Annotations in red or orange correspond to the presence or absence of genes encoding predicted secreted lipase/esterase and peptidase/proteases, respectively (see Supplementary Table 6 for annotations and sequences). Bootstrap values of >90% are indicated by filled black circles. The scale bar represents 10% sequence divergence.

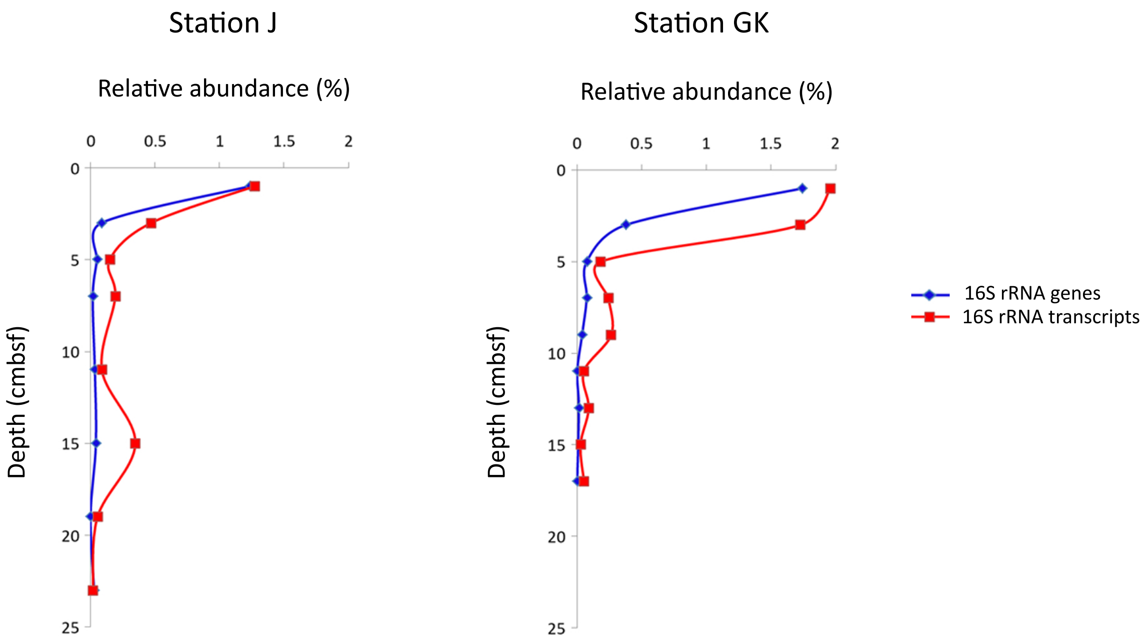


**Supplementary Figure S6**. Relative abundances of *Psychromonas* spp. in sediments of Smeerenbergfjord, Svalbard, as determined by 16S rRNA-gene and -transcript (cDNA) amplicon sequencing. Cmbsf = Centimeters below seafloor.

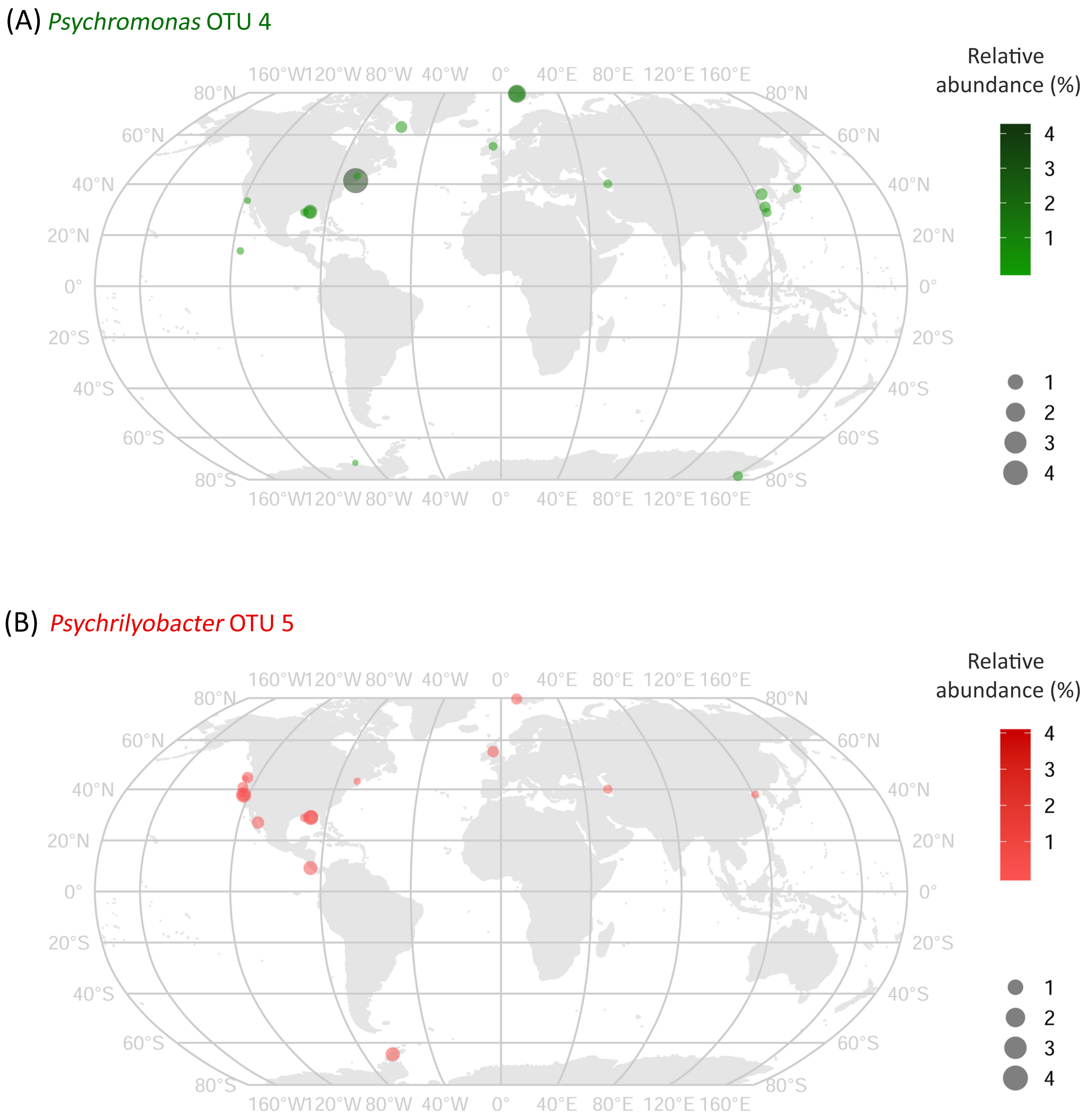


**Supplementary Figure S7**. Presence and relative abundances of *Psychromonas* OTU 4 (A) *Psychrilyobacter* OTU 5 (B) related sequences (>97%) in publically available 16S rRNA gene datasets, as determined by IMNGS analyses (see Materials and Methods). Additional samples from Svalbard that were determined in this study, are also included.

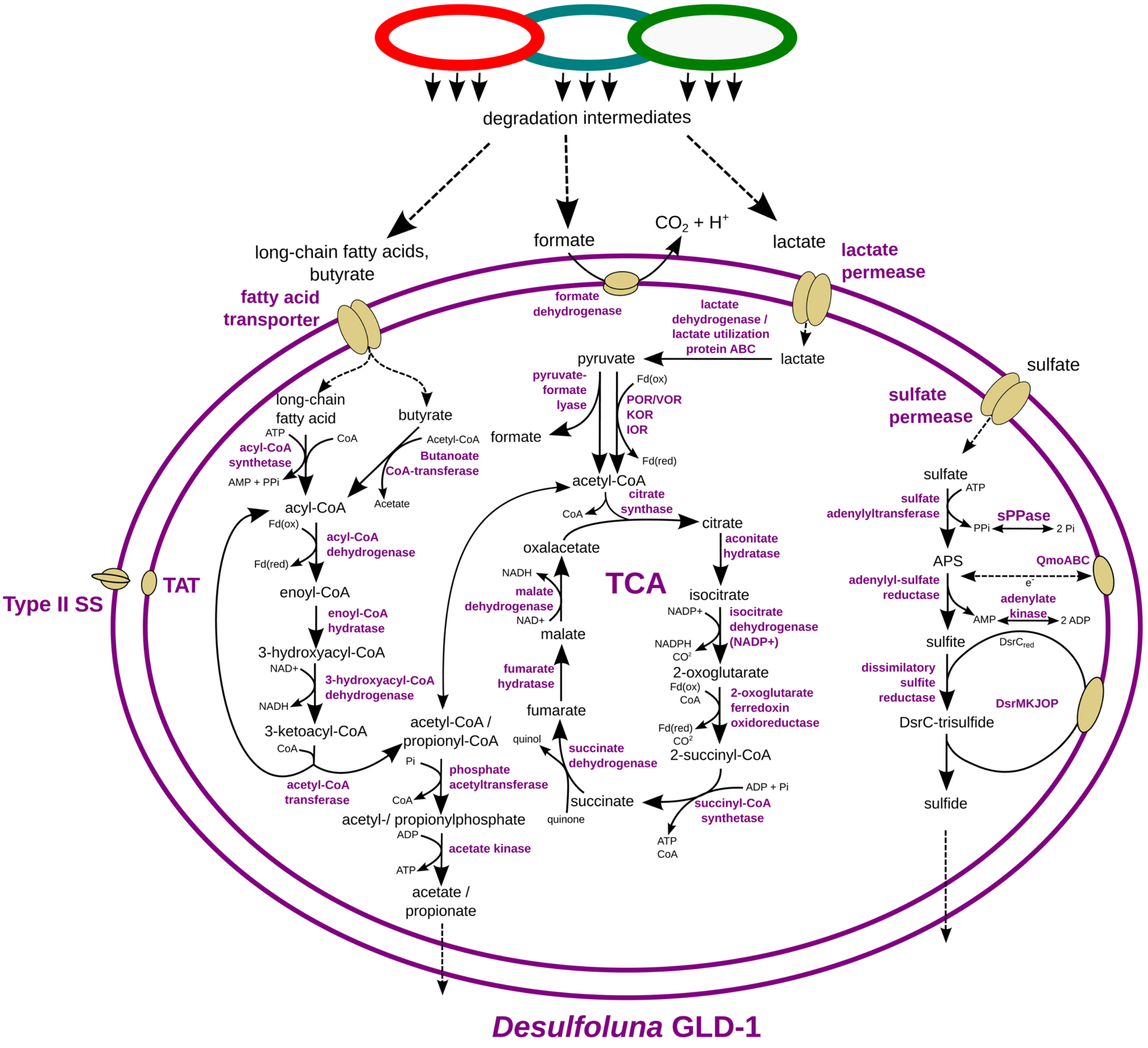


**Supplementary Figure 8**. Genome-inferred metabolic model of *Desulfoluna* MAG GLD-1. SS, Secretion system; TCA, tricarboxylic acid cycle; TAT, Twin-arginine translocation system.
