## Supplementary Tables 1-6 for "Anaerobic microbial degradation of protein and lipid macromolecules in subarctic marine sediment"

**Supplementary Table S1. Summary of MEROPS and ESTHER database description for peptidases and lipases, respectively.**

| | **Peptidases** | |
|---|---|---|
| **MEROPS ID** | **MEROPS description summary** | **Selected for further analyses** |
| A8 | production of the bacterial cell wall | No |
| M1 | some are cleaving amino acids from small peptides | Yes |
| M3 | Lactobacillus oligoendopeptidase cleaves oligopeptides | Yes |
| M14 | digestion of food and other functions | Yes |
| M20 | conversion of proteins to free amino acids. | Yes |
| M24 | removes N-terminal Methionine from protein | Yes |
| M28 | alkaline phosphatase isozyme conversion | No |
| M29 | catabolic peptidase in Streptococcus thermophilus | Yes |
| M42 | used by Lactobacillus for nutrition on milk-protein powder | Yes |
| M48 | degradation of abnormal proteins | No |
| M50 | regulated intermembrane proetolysis | No |
| S8 | probably involved in nutrition | Yes |
| S12 | synthesis and remodelling of bacterial cell walls | No |
| S15 | degradation of casein | Yes |
| S24 | regulatory stress response | No |
| S26 | remove the signal peptides and facilitate secretion | No |

| | **Lipases** | |
|---|---|---|
| **ESTHER family** | **ESTHER description summary** | **Selected for further analyses** |
| 6_AlphaBeta_hydrolase | not well characterized | No |
| Abhydrolase_7 | acetyl-esterase_deacetylase or Dienelactone_hydrolase or xylan esterase | No |
| Bacterial_EstLip_FamX | Family of Bacterial lipolytic enzymes | Yes |
| Carb_B_Bacteria | carboxylesterase, type B | Yes |
| CarbLipBact_2 | members of this group are esterases, lipases | Yes |
| Duf_1023 | proteins of unknown function | No |
| Duf_3089 | lipolytic enzyme family defined from isolation and characterization of two esterases from a metagenomic library | Yes |
| Duf_900 | proteins of unknown function | No |
| Epoxide_hydrolase | conversion of epoxides to corresponding diols | No |
| GTSAGmotif | lipases from metagenomic library | Yes |
| Hormone-sensitive_lipase_like | esterase/lipase, family IV | Yes |
| Lipase_2 | lipases | Yes |
| Lipase_3 | lipases | Yes |
| Monoglyceridelipase_lysophospholip | has some homology to peptidases S33 | No |
| NFM-deformylase | N-formylmaleamic acid to formic and maleamic acid | No |
| PGAP1 | attachment to proteins factor 1 | No |

**Supplementary Table S2. Statistics results**

| Day | Comparison | Compound | p_value | adjusted_p_value |
|---|---|---|---|---|
| Day_0 | Proteins vs No_Substrate | Formate_µM | 1.000 | 1.000 |
| Day_2 | Proteins vs No_Substrate | Formate_µM | 0.043 | 0.107 |
| Day_5 | Proteins vs No_Substrate | Formate_µM | 0.483 | 0.594 |
| Day_10 | Proteins vs No_Substrate | Formate_µM | 0.401 | 0.515 |
| Day_17 | Proteins vs No_Substrate | Formate_µM | 0.247 | 0.349 |
| Day_0 | Proteins vs No_Substrate | Acetate_µM | 1.000 | 1.000 |
| Day_2 | Proteins vs No_Substrate | Acetate_µM | 0.009 | 0.035 |
| Day_5 | Proteins vs No_Substrate | Acetate_µM | 0.005 | 0.022 |
| Day_10 | Proteins vs No_Substrate | Acetate_µM | 0.036 | 0.094 |
| Day_17 | Proteins vs No_Substrate | Acetate_µM | 0.097 | 0.185 |
| Day_0 | Proteins vs No_Substrate | Propionate_µM | 1.000 | 1.000 |
| Day_2 | Proteins vs No_Substrate | Propionate_µM | 0.010 | 0.036 |
| Day_5 | Proteins vs No_Substrate | Propionate_µM | 0.000 | 0.002 |
| Day_10 | Proteins vs No_Substrate | Propionate_µM | 0.223 | 0.321 |
| Day_17 | Proteins vs No_Substrate | Propionate_µM | 0.018 | 0.054 |
| Day_0 | Proteins vs No_Substrate | Butyrate_µM | NaN | NaN |
| Day_2 | Proteins vs No_Substrate | Butyrate_µM | 0.010 | 0.035 |
| Day_5 | Proteins vs No_Substrate | Butyrate_µM | 0.003 | 0.018 |
| Day_10 | Proteins vs No_Substrate | Butyrate_µM | 0.598 | 0.721 |
| Day_17 | Proteins vs No_Substrate | Butyrate_µM | 0.137 | 0.232 |
| Day_0 | Proteins vs No_Substrate | iso.Butyrate_µM | NaN | NaN |
| Day_2 | Proteins vs No_Substrate | iso.Butyrate_µM | 0.003 | 0.017 |
| Day_5 | Proteins vs No_Substrate | iso.Butyrate_µM | 0.005 | 0.022 |
| Day_10 | Proteins vs No_Substrate | iso.Butyrate_µM | 0.000 | 0.004 |
| Day_17 | Proteins vs No_Substrate | iso.Butyrate_µM | 0.067 | 0.146 |
| Day_0 | Proteins vs No_Substrate | Lactate_µM | NaN | NaN |
| Day_2 | Proteins vs No_Substrate | Lactate_µM | NaN | NaN |
| Day_5 | Proteins vs No_Substrate | Lactate_µM | NaN | NaN |
| Day_10 | Proteins vs No_Substrate | Lactate_µM | NaN | NaN |
| Day_17 | Proteins vs No_Substrate | Lactate_µM | NaN | NaN |
| Day_0 | Proteins vs Proteins_inhibited | Formate_µM | 1.000 | 1.000 |
| Day_2 | Proteins vs Proteins_inhibited | Formate_µM | 0.341 | 0.448 |
| Day_5 | Proteins vs Proteins_inhibited | Formate_µM | 0.161 | 0.258 |
| Day_10 | Proteins vs Proteins_inhibited | Formate_µM | 0.191 | 0.283 |
| Day_17 | Proteins vs Proteins_inhibited | Formate_µM | 0.440 | 0.556 |
| Day_0 | Proteins vs Proteins_inhibited | Acetate_µM | 1.000 | 1.000 |
| Day_2 | Proteins vs Proteins_inhibited | Acetate_µM | 0.132 | 0.225 |
| Day_5 | Proteins vs Proteins_inhibited | Acetate_µM | 0.001 | 0.010 |
| Day_10 | Proteins vs Proteins_inhibited | Acetate_µM | 0.000 | 0.004 |
| Day_17 | Proteins vs Proteins_inhibited | Acetate_µM | 0.015 | 0.047 |
| Day_0 | Proteins vs Proteins_inhibited | Propionate_µM | 1.000 | 1.000 |
| Day_2 | Proteins vs Proteins_inhibited | Propionate_µM | 0.005 | 0.022 |
| Day_5 | Proteins vs Proteins_inhibited | Propionate_µM | 0.001 | 0.010 |
| Day_10 | Proteins vs Proteins_inhibited | Propionate_µM | 0.005 | 0.022 |
| Day_17 | Proteins vs Proteins_inhibited | Propionate_µM | 0.001 | 0.010 |
| Day_0 | Proteins vs Proteins_inhibited | Butyrate_µM | NaN | NaN |
| Day_2 | Proteins vs Proteins_inhibited | Butyrate_µM | 0.628 | 0.741 |
| Day_5 | Proteins vs Proteins_inhibited | Butyrate_µM | 0.044 | 0.107 |
| Day_10 | Proteins vs Proteins_inhibited | Butyrate_µM | 0.047 | 0.110 |
| Day_17 | Proteins vs Proteins_inhibited | Butyrate_µM | 0.098 | 0.185 |
| Day_0 | Proteins vs Proteins_inhibited | iso.Butyrate_µM | NaN | NaN |
| Day_2 | Proteins vs Proteins_inhibited | iso.Butyrate_µM | 0.032 | 0.090 |
| Day_5 | Proteins vs Proteins_inhibited | iso.Butyrate_µM | 0.085 | 0.173 |
| Day_10 | Proteins vs Proteins_inhibited | iso.Butyrate_µM | 0.851 | 0.980 |
| Day_17 | Proteins vs Proteins_inhibited | iso.Butyrate_µM | 0.162 | 0.258 |
| Day_0 | Proteins vs Proteins_inhibited | Lactate_µM | NaN | NaN |
| Day_2 | Proteins vs Proteins_inhibited | Lactate_µM | NaN | NaN |
| Day_5 | Proteins vs Proteins_inhibited | Lactate_µM | NaN | NaN |
| Day_10 | Proteins vs Proteins_inhibited | Lactate_µM | NaN | NaN |

| | | | | |
|---|---|---|---|---|
| Day_17 | Proteins vs Proteins_inhibited | Lactate_µM | NaN | NaN |
| Day_0 | Proteins_inhibited vs No Substrate | Formate_µM | 1.000 | 1.000 |
| Day_2 | Proteins_inhibited vs No Substrate | Formate_µM | 0.183 | 0.276 |
| Day_5 | Proteins_inhibited vs No Substrate | Formate_µM | 0.157 | 0.258 |
| Day_10 | Proteins_inhibited vs No Substrate | Formate_µM | 0.200 | 0.293 |
| Day_17 | Proteins_inhibited vs No Substrate | Formate_µM | 0.174 | 0.272 |
| Day_0 | Proteins_inhibited vs No Substrate | Acetate_µM | 1.000 | 1.000 |
| Day_2 | Proteins_inhibited vs No Substrate | Acetate_µM | 0.481 | 0.594 |
| Day_5 | Proteins_inhibited vs No Substrate | Acetate_µM | 0.037 | 0.096 |
| Day_10 | Proteins_inhibited vs No Substrate | Acetate_µM | 0.113 | 0.209 |
| Day_17 | Proteins_inhibited vs No Substrate | Acetate_µM | 0.083 | 0.173 |
| Day_0 | Proteins_inhibited vs No Substrate | Propionate_µM | 1.000 | 1.000 |
| Day_2 | Proteins_inhibited vs No Substrate | Propionate_µM | 0.601 | 0.721 |
| Day_5 | Proteins_inhibited vs No Substrate | Propionate_µM | 0.052 | 0.117 |
| Day_10 | Proteins_inhibited vs No Substrate | Propionate_µM | 0.393 | 0.509 |
| Day_17 | Proteins_inhibited vs No Substrate | Propionate_µM | 0.109 | 0.204 |
| Day_0 | Proteins_inhibited vs No Substrate | Butyrate_µM | NaN | NaN |
| Day_2 | Proteins_inhibited vs No Substrate | Butyrate_µM | 0.049 | 0.115 |
| Day_5 | Proteins_inhibited vs No Substrate | Butyrate_µM | 0.036 | 0.094 |
| Day_10 | Proteins_inhibited vs No Substrate | Butyrate_µM | 0.046 | 0.110 |
| Day_17 | Proteins_inhibited vs No Substrate | Butyrate_µM | 0.056 | 0.124 |
| Day_0 | Proteins_inhibited vs No Substrate | iso.Butyrate_µM | NaN | NaN |
| Day_2 | Proteins_inhibited vs No Substrate | iso.Butyrate_µM | 0.084 | 0.173 |
| Day_5 | Proteins_inhibited vs No Substrate | iso.Butyrate_µM | 0.028 | 0.081 |
| Day_10 | Proteins_inhibited vs No Substrate | iso.Butyrate_µM | 0.087 | 0.173 |
| Day_17 | Proteins_inhibited vs No Substrate | iso.Butyrate_µM | 0.089 | 0.173 |
| Day_0 | Proteins_inhibited vs No Substrate | Lactate_µM | NaN | NaN |
| Day_2 | Proteins_inhibited vs No Substrate | Lactate_µM | NaN | NaN |
| Day_5 | Proteins_inhibited vs No Substrate | Lactate_µM | NaN | NaN |
| Day_10 | Proteins_inhibited vs No Substrate | Lactate_µM | NaN | NaN |
| Day_17 | Proteins_inhibited vs No Substrate | Lactate_µM | NaN | NaN |
| Day_0 | Lipids vs No_Substrate | Formate_µM | 1.000 | 1.000 |
| Day_2 | Lipids vs No_Substrate | Formate_µM | 0.163 | 0.258 |
| Day_5 | Lipids vs No_Substrate | Formate_µM | 0.271 | 0.372 |
| Day_10 | Lipids vs No_Substrate | Formate_µM | 0.342 | 0.448 |
| Day_17 | Lipids vs No_Substrate | Formate_µM | 0.284 | 0.382 |
| Day_0 | Lipids vs No_Substrate | Acetate_µM | 1.000 | 1.000 |
| Day_2 | Lipids vs No_Substrate | Acetate_µM | 0.119 | 0.211 |
| Day_5 | Lipids vs No_Substrate | Acetate_µM | 0.127 | 0.220 |
| Day_10 | Lipids vs No_Substrate | Acetate_µM | 0.114 | 0.209 |
| Day_17 | Lipids vs No_Substrate | Acetate_µM | 0.444 | 0.556 |
| Day_0 | Lipids vs No_Substrate | Propionate_µM | 1.000 | 1.000 |
| Day_2 | Lipids vs No_Substrate | Propionate_µM | 0.036 | 0.094 |
| Day_5 | Lipids vs No_Substrate | Propionate_µM | 0.159 | 0.258 |
| Day_10 | Lipids vs No_Substrate | Propionate_µM | 0.270 | 0.372 |
| Day_17 | Lipids vs No_Substrate | Propionate_µM | 0.085 | 0.173 |
| Day_0 | Lipids vs No_Substrate | Butyrate_µM | NaN | NaN |
| Day_2 | Lipids vs No_Substrate | Butyrate_µM | 0.042 | 0.107 |
| Day_5 | Lipids vs No_Substrate | Butyrate_µM | 0.002 | 0.014 |
| Day_10 | Lipids vs No_Substrate | Butyrate_µM | 0.016 | 0.051 |
| Day_17 | Lipids vs No_Substrate | Butyrate_µM | 0.018 | 0.054 |
| Day_0 | Lipids vs No_Substrate | iso.Butyrate_µM | NaN | NaN |
| Day_2 | Lipids vs No_Substrate | iso.Butyrate_µM | 0.017 | 0.053 |
| Day_5 | Lipids vs No_Substrate | iso.Butyrate_µM | 0.013 | 0.044 |
| Day_10 | Lipids vs No_Substrate | iso.Butyrate_µM | 0.002 | 0.015 |
| Day_17 | Lipids vs No_Substrate | iso.Butyrate_µM | 0.008 | 0.033 |
| Day_0 | Lipids vs No_Substrate | Lactate_µM | NaN | NaN |
| Day_2 | Lipids vs No_Substrate | Lactate_µM | NaN | NaN |
| Day_5 | Lipids vs No_Substrate | Lactate_µM | NaN | NaN |
| Day_10 | Lipids vs No_Substrate | Lactate_µM | NaN | NaN |
| Day_17 | Lipids vs No_Substrate | Lactate_µM | NaN | NaN |
| Day_0 | Lipids vs Lipids_inhibited | Formate_µM | 1.000 | 1.000 |

| | | | | |
|---|---|---|---|---|
| Day_2 | Lipids vs Lipids_inhibited | Formate_µM | 0.235 | 0.335 |
| Day_5 | Lipids vs Lipids_inhibited | Formate_µM | 0.001 | 0.010 |
| Day_10 | Lipids vs Lipids_inhibited | Formate_µM | 0.003 | 0.018 |
| Day_17 | Lipids vs Lipids_inhibited | Formate_µM | 0.000 | 0.004 |
| Day_0 | Lipids vs Lipids_inhibited | Acetate_µM | 1.000 | 1.000 |
| Day_2 | Lipids vs Lipids_inhibited | Acetate_µM | 0.119 | 0.211 |
| Day_5 | Lipids vs Lipids_inhibited | Acetate_µM | 0.074 | 0.159 |
| Day_10 | Lipids vs Lipids_inhibited | Acetate_µM | 0.000 | 0.002 |
| Day_17 | Lipids vs Lipids_inhibited | Acetate_µM | 0.011 | 0.036 |
| Day_0 | Lipids vs Lipids_inhibited | Propionate_µM | 1.000 | 1.000 |
| Day_2 | Lipids vs Lipids_inhibited | Propionate_µM | 0.259 | 0.362 |
| Day_5 | Lipids vs Lipids_inhibited | Propionate_µM | 0.295 | 0.394 |
| Day_10 | Lipids vs Lipids_inhibited | Propionate_µM | 0.052 | 0.117 |
| Day_17 | Lipids vs Lipids_inhibited | Propionate_µM | 0.120 | 0.211 |
| Day_0 | Lipids vs Lipids_inhibited | Butyrate_µM | NaN | NaN |
| Day_2 | Lipids vs Lipids_inhibited | Butyrate_µM | 0.994 | 1.000 |
| Day_5 | Lipids vs Lipids_inhibited | Butyrate_µM | 0.001 | 0.010 |
| Day_10 | Lipids vs Lipids_inhibited | Butyrate_µM | 0.002 | 0.015 |
| Day_17 | Lipids vs Lipids_inhibited | Butyrate_µM | 0.001 | 0.010 |
| Day_0 | Lipids vs Lipids_inhibited | iso.Butyrate_µM | NaN | NaN |
| Day_2 | Lipids vs Lipids_inhibited | iso.Butyrate_µM | 0.223 | 0.321 |
| Day_5 | Lipids vs Lipids_inhibited | iso.Butyrate_µM | 0.009 | 0.034 |
| Day_10 | Lipids vs Lipids_inhibited | iso.Butyrate_µM | 0.001 | 0.010 |
| Day_17 | Lipids vs Lipids_inhibited | iso.Butyrate_µM | 0.013 | 0.042 |
| Day_0 | Lipids vs Lipids_inhibited | Lactate_µM | NaN | NaN |
| Day_2 | Lipids vs Lipids_inhibited | Lactate_µM | 0.184 | 0.276 |
| Day_5 | Lipids vs Lipids_inhibited | Lactate_µM | 0.004 | 0.020 |
| Day_10 | Lipids vs Lipids_inhibited | Lactate_µM | 0.004 | 0.021 |
| Day_17 | Lipids vs Lipids_inhibited | Lactate_µM | NaN | NaN |
| Day_0 | Lipids_inhibited vs No Substrate | Formate_µM | 1.000 | 1.000 |
| Day_2 | Lipids_inhibited vs No Substrate | Formate_µM | 0.184 | 0.276 |
| Day_5 | Lipids_inhibited vs No Substrate | Formate_µM | 0.001 | 0.010 |
| Day_10 | Lipids_inhibited vs No Substrate | Formate_µM | 0.003 | 0.018 |
| Day_17 | Lipids_inhibited vs No Substrate | Formate_µM | 0.001 | 0.010 |
| Day_0 | Lipids_inhibited vs No Substrate | Acetate_µM | 1.000 | 1.000 |
| Day_2 | Lipids_inhibited vs No Substrate | Acetate_µM | 0.436 | 0.556 |
| Day_5 | Lipids_inhibited vs No Substrate | Acetate_µM | 0.033 | 0.091 |
| Day_10 | Lipids_inhibited vs No Substrate | Acetate_µM | 0.163 | 0.258 |
| Day_17 | Lipids_inhibited vs No Substrate | Acetate_µM | 0.089 | 0.173 |
| Day_0 | Lipids_inhibited vs No Substrate | Propionate_µM | 1.000 | 1.000 |
| Day_2 | Lipids_inhibited vs No Substrate | Propionate_µM | 0.635 | 0.744 |
| Day_5 | Lipids_inhibited vs No Substrate | Propionate_µM | 0.000 | 0.004 |
| Day_10 | Lipids_inhibited vs No Substrate | Propionate_µM | 0.720 | 0.836 |
| Day_17 | Lipids_inhibited vs No Substrate | Propionate_µM | 0.533 | 0.650 |
| Day_0 | Lipids_inhibited vs No Substrate | Butyrate_µM | NaN | NaN |
| Day_2 | Lipids_inhibited vs No Substrate | Butyrate_µM | 0.279 | 0.380 |
| Day_5 | Lipids_inhibited vs No Substrate | Butyrate_µM | 0.001 | 0.010 |
| Day_10 | Lipids_inhibited vs No Substrate | Butyrate_µM | 0.002 | 0.016 |
| Day_17 | Lipids_inhibited vs No Substrate | Butyrate_µM | 0.002 | 0.016 |
| Day_0 | Lipids_inhibited vs No Substrate | iso.Butyrate_µM | NaN | NaN |
| Day_2 | Lipids_inhibited vs No Substrate | iso.Butyrate_µM | 0.610 | 0.726 |
| Day_5 | Lipids_inhibited vs No Substrate | iso.Butyrate_µM | 0.007 | 0.029 |
| Day_10 | Lipids_inhibited vs No Substrate | iso.Butyrate_µM | 0.000 | 0.007 |
| Day_17 | Lipids_inhibited vs No Substrate | iso.Butyrate_µM | 0.009 | 0.033 |
| Day_0 | Lipids_inhibited vs No Substrate | Lactate_µM | NaN | NaN |
| Day_2 | Lipids_inhibited vs No Substrate | Lactate_µM | 0.184 | 0.276 |
| Day_5 | Lipids_inhibited vs No Substrate | Lactate_µM | 0.004 | 0.020 |
| Day_10 | Lipids_inhibited vs No Substrate | Lactate_µM | 0.004 | 0.021 |
| Day_17 | Lipids_inhibited vs No Substrate | Lactate_µM | NaN | NaN |

**Supplementary Table S3. Significant enrichment of 16S rRNA gene OTUs in $^{13}C$ fractions of DNA stable isotope gradients.**

| OTU | Taxonomy | base mean | log2-fold change | adjusted p-value | Treatment | Day |
|---|---|---|---|---|---|---|
| OTU_80 | *Gammaproteobacteria* , *Vibrionaceae* | 8.7 | 7.7 | 0.000 | Proteins | 10 |
| OTU_80 | Gammaproteobacteria, Vibrionaceae | 8.7 | 6.7 | 0.002 | Lipids | 5 |
| OTU_232 | *Deltaproteobacteria* , *Desulfatiglans* | 2.4 | 6.4 | 0.005 | Proteins | 10 |
| OTU_4 | *Gammaproteobacteria* , *Psychromonas* | 936.5 | 5.5 | 0.000 | Lipids | 5 |
| OTU_312 | *Cyanobacteria* | 1.9 | 5.2 | 0.034 | Proteins | 10 |
| OTU_4 | *Gammaproteobacteria* , *Psychromonas* | 936.5 | 4.9 | 0.000 | Lipids | 10 |
| OTU_4141 | *Firmicutes* , *Defluviitaleaceae* | 5.2 | 4.8 | 0.015 | Lipids | 10 |
| OTU_183 | *Deltaproteobacteria* , Sva0485 | 4.2 | 4.8 | 0.003 | Proteins | 10 |
| OTU_184 | *Bacteroidetes* , *Prolixibacter* | 1.4 | 4.5 | 0.083 | Proteins | 10 |
| OTU_123 | *Marinimicrobia* | 4.3 | 4.5 | 0.007 | Proteins | 10 |
| OTU_4 | *Gammaproteobacteria* , *Psychromonas* | 936.5 | 4.5 | 0.000 | Proteins | 5 |
| OTU_80 | *Gammaproteobacteria* , *Vibrionaceae* | 8.7 | 4.3 | 0.015 | Lipids | 10 |
| OTU_124 | *Bacteroidetes* , BD2-2 | 3.6 | 3.7 | 0.055 | Proteins | 10 |
| OTU_285 | *Gammaproteobacteria* , JTB255 | 2.7 | 3.6 | 0.083 | Proteins | 10 |
| OTU_13310 | *Deltaproteobacteria* , *Desulfofrigus* | 5.4 | 3.5 | 0.072 | Lipids | 10 |
| OTU_4719 | *Deltaproteobacteria* , NB1-J | 3.1 | 3.3 | 0.098 | Proteins | 10 |
| OTU_54 | *Gammaproteobacteria* , *Vibrionaceae* | 9.1 | 3.2 | 0.030 | Proteins | 5 |
| OTU_128 | *Gammaproteobacteria* , Sva0071 | 4.1 | 3.1 | 0.045 | Proteins | 10 |
| OTU_1 | *Firmicutes* , JTB215 | 967.1 | 3.0 | 0.018 | Proteins | 5 |
| OTU_202 | *Firmicutes* , *Clostridiales* | 2.1 | 3.0 | 0.098 | Proteins | 10 |
| OTU_205 | *Gammaproteobacteria* , Thiotrichaceae | 3.2 | 3.0 | 0.083 | Proteins | 10 |
| OTU_4050 | *Firmicutes* , *Fusibacter* | 25.8 | 3.0 | 0.001 | Proteins | 10 |
| OTU_4 | *Gammaproteobacteria* , *Psychromonas* | 936.5 | 2.8 | 0.009 | Proteins | 10 |
| OTU_1 | *Firmicutes* , JTB215 | 967.1 | 2.8 | 0.036 | Lipids | 10 |
| OTU_38 | *Firmicutes* , *Clostridiales* | 11.2 | 2.6 | 0.006 | Proteins | 5 |
| OTU_19 | *Deltaproteobacteria* , SEEP-SRB1 | 22.5 | 2.1 | 0.009 | Proteins | 10 |
| OTU_892 | *Deltaproteobacteria* , *Desulfobulbaceae* | 11.0 | 2.1 | 0.098 | Proteins | 10 |
| OTU_5 | *Fusobacteria* , *Psychrilyobacter* | 297.5 | 2.0 | 0.005 | Proteins | 5 |
| OTU_36 | *Deltaproteobacteria* , *Desulfobulbaceae* | 16.0 | 2.0 | 0.068 | Proteins | 10 |
| OTU_44 | *Deltaproteobacteria* , *Desulfatiglans* | 16.5 | 1.8 | 0.098 | Proteins | 10 |
| OTU_67 | *Deltaproteobacteria* , *Desulfoconvexum* | 19.0 | 1.7 | 0.055 | Proteins | 10 |
| OTU_2 | *Deltaproteobacteria* , *Desulfofrigus* | 699.6 | 1.3 | 0.043 | Proteins | 10 |
| OTU_5 | *Fusobacteria* , *Psychrilyobacter* | 297.5 | 1.3 | 0.055 | Proteins | 10 |

**Supplementary Table S4. Genome statistics of metagenome-assembled genomes.**

| Bin | Genbank accession no. | Microscope annotation ID | GTDB classification | Closest relative [ANI (% aligned) / AAI (% aligned)] | Completeness | Contamination | Strain heterogeneity |
|---|---|---|---|---|---|---|---|
| *Psychromonas* GLG-1 | SAMN14421524 | 42481_Bin_4 | d__Bacteria;p__Proteobacteria;c__Gammaproteobacteria;o__Enterobacterales;f__Psychromonadaceae;g__Psychromonas;s_ | *Psychromonas aquimarina* ATCC BAA-1526 GCF_000381745.1 [80.7 (41.8) / 80.73 (66.28)] | 97.39 | 0.81 | 50 |
| *Clostridia* GPF-1 | SAMN14421525 | 42482_Bin_3 | d__Bacteria;p__Firmicutes_A;c__Clostridia;o__Tissierellales;f__;g__;s__ | *Caloranaerobacter azorensis* DSM 13643 GCA_900129995.1 [ - / 53.39 (37)] | 99.07 | 0.7 | 0 |
| *Desulfoluna* GLD-1 | SAMN14421526 | 42483_Bin_2 | d__Bacteria;p__Desulfobacterota;c__Desulfobacteria;o__Desulfobacterales;f__Desulfobacteraceae;g__Desulfoluna;s__ | *Desulfoluna spongiiphila* GCA_900101345.1 [79.42 (39.41) / 76.93 (68.72)] | 94.95 | 1.36 | 0 |
| 42485_Bin_1 | SAMN14421527 | . | d__Bacteria;p__Desulfuromonadota;c__Desulfuromonadia;o__Desulfuromonadales;f__Pelobacteraceae_A;g__SFB93;s__ | *Pelobacter* sp. SFB93 GCF_001887775.1 [80.99 (60.65) / 84.98 (67.80)] | 98.71 | 1.47 | 14.29 |
| 42482_Bin_1 | SAMN14421528 | . | d__Bacteria;p__Desulfobacterota;c__Syntrophobacteria;o__BM002;f__BM002;g__BM002;s__BM002 sp002899795 | *Desulfobacteraceae* bacterium GCA_002899795.1 [95.31 (73.68) / 95.26 (64.16)] | 96.61 | 3.39 | 0 |
| 40935_Bin_6_Spades | SAMN14421529 | . | d__Bacteria;p__Desulfobacterota;c__Desulfobacteria;o__Desulfobacterales;f__BuS5;g__;s__ | *Desulfobacteraceae* bacterium GCA_002868985.1 [79.31 (38.02) / 76.48 (45.11)] | 87.49 | 2.71 | 30 |
| 42483_Bin_0_Spades_Anvio | SAMN14421530 | . | d__Bacteria;p__Firmicutes_A;c__Clostridia;o__Peptostreptococcales;f__;g__;s__ | *Caminicella sporogenes* DSM 14501 GCF_900142285.1 [ - / 56.06 (42.81)] | 85.99 | 4.25 | 0 |
| 40935_Bin_7_Spades | SAMN14421531 | . | d__Archaea;p__Crenarchaeota;c__Bathyarchaeia;o__TCS64;f__TCS64;g__RBG-16-57-9;s__ | Candidatus *Bathyarchaeota* archaeon GCA_004525915.1 [ 77.69 (15.99) / 71.11 (51.03)] | 83.64 | 3.74 | 0 |
| 42485_Bin_1_Spades | SAMN14421532 | . | d__Bacteria;p__Proteobacteria;c__Alphaproteobacteria;o__Micavibrionales;f__;g__;s__ | *Azospirillum* sp. K2W22B-5 GCF_003590795.1 [ - / 55.67 (43.89)] | 80.7 | 0.43 | 0 |

**Supplementary Table S5. Annotation of MAGs and the genome of *Psychrilyobacter atlanticus*.**

| | *Psychrilyobacter atlanticus* | *Psychromonas* GLG-1 | *Clostridiales* GPF-1 | *Desulfoluna* GLD-1 |
|---|---|---|---|---|
| **Extracellular peptidases** | | | | |
| M3 | K337_v1_10834 | - | - | - |
| M20 | - | - | FUSI_v1_490026 | - |
| M24 | K337_v1_10582 | - | FUSI_v1_750006 | - |
| S8 | - | - | FUSI_v1_960008 | - |
| **Intracellular peptidases** | | | | |
| M1 | K337_v1_20320 | PSYM_v1_1690003 | - | DESP_v1_990015 |
| M3 | K337_v1_10060; K337_v1_10959 | PSYM_v1_200017 | FUSI_v1_620036 | DESP_v1_1000010 |
| M14 | - | - | FUSI_v1_430016 | - |
| M20 | ; K337_v1_20729; K337_v1_20883; | M_v1_1880004; PSYM_v1_77 | v1_520043; FUSI_v1_6900 | SP_v1_450039; DESP_v1_95 |
| M24 | ; K337_v1_30031; K337_v1_11495; | YM_v1_40024; PSYM_v1_190 | _v1_640120; FUSI_v1_94 | P_v1_210001; DESP_v1_670 |
| M29 | K337_v1_10108 | - | FUSI_v1_670015 | - |
| M42 | ; K337_v1_20380; K337_v1_20685; | - | 071; FUSI_v1_640072; FU | - |
| S15 | - | PSYM_v1_250021 | - | DESP_v1_550012 |
| **Extracellular lipases (ESTHER classification)** | | | | |
| Carboxylesterase, type B (carboxylesterase) | - | PSYM_v1_1870005 | - | - |
| Bacterial_EstLip_FamX (esterase / lipase) | - | PSYM_v1_540010 | FUSI_v1_10055 | - |
| **Membrane lipases** | | | | |
| Carboxylesterase, type B (carboxylesterase) | - | - | FUSI_v1_1090001 | - |
| Bacterial_EstLip_FamX (esterase / lipase) | - | - | FUSI_v1_840070 | - |
| **Intracellular lipases** | | | | |
| Carboxylesterase, type B (carboxylesterase) | - | PSYM_v1_2290005 | | DESP_v1_250018 |
| Hormone-sensitive_lipase_like (hormone-sensitive lipase) | - | - | _v1_1110043; FUSI_v1_12 | - |
| Lipase_3 (lipase) | - | - | - | DESP_v1_1460001 |
| CarbLipBact_2 (carboxylesterase) | - | - | - | P_v1_330004; DESP_v1_170 |
| **Extracellular glycoside hydrolases** | | | | |
| GH 1 | - | M_v1_2060004; PSYM_v1_54 | - | - |
| GH 3 | K337_v1_11729 | _v1_1230007; PSYM_v1_287 | - | - |
| GH 5 | - | - | - | - |
| GH 9 | - | - | - | DESP_v1_770006 |
| GH 13 | - | _v1_1260002; PSYM_v1_140 | FUSI_v1_1250081 | - |
| GH 15 | - | - | - | - |
| GH 16 | - | PSYM_v1_700008 | - | - |
| GH 17 | - | PSYM_v1_700006 | - | - |
| GH 18 | - | PSYM_v1_1680004 | - | - |
| GH 20 | - | - | - | - |
| GH 26 | - | - | FUSI_v1_320001 | - |
| GH 31 | - | - | - | - |
| GH 81 | - | PSYM_v1_700005 | - | - |
| **Peptide, amino acid transporters** | | | | |
| peptide ABC transporter solute-binding protein | 1_21084; K337_v1_10066; K337_v1 | 12; PSYM_v1_1890004; PSY | 9; FUSI_v1_430021; FUSI_7 | ; DESP_v1_1230003; DESP_ |
| peptide ABC transporter permease | ; K337_v1_21082; K337_v1_21083; | _v1_1890002; PSYM_v1_189 | _v1_490040; FUSI_v1_490 | 5; DESP_v1_190003; DESP_ |
| peptide ABC transporter ATP-binding protein | ; K337_v1_21080; K337_v1_21081; | 11; PSYM_v1_1890001; PSY | _v1_490038; FUSI_v1_490 | 006; DESP_v1_190006; DES |
| proton-dependent oligopeptide transporter | ; K337_v1_11558; K337_v1_20326; | - | - | - |
| oligopeptide transporter, OPT superfamily | K337_v1_12103 | - | - | - |
| (branched-chain) amino acid ABC transporter substrate-binding protein | 1_20143; K337_v1_10045; K337_v1 | 010; PSYM_v1_1780001; PSY | 3; FUSI_v1_560002; FUSI_ | SP_v1_990022; DESP_v1_14 |
| (branched-chain) amino acid ABC transporter permease | 1_10042; K337_v1_10043; K337_v1 | 002; PSYM_v1_1780003; PS | v1_450021; FUSI_v1_5600 | SP_v1_1440006; DESP_v1_1 |
| (branched-chain) amino acid ABC transporter ATP-binding protein | 1; K337_v1_10044; K337_v1_10748; | _v1_3310003; PSYM_v1_22 | 2; FUSI_v1_560005; FUSI_ | SP_v1_130004; DESP_v1_17 |
| amino acid transporter (not ABC-type) | 2; K337_v1_11837; K337_v1_11930; | 006; PSYM_v1_1000001; PS | v1_430065; FUSI_v1_5400 | 024; DESP_v1_140024; DES |
| **Fatty acid transporters** | | | | |
| short-chain fatty acid transporter | K337_v1_10807 | - | - | - |
| long-chain fatty acid transporter | - | M_v1_3160003; PSYM_v1_24 | - | DESP_v1_1780005 |
| lipid carrier protein | - | PSYM_v1_1250003 | - | - |
| **Glutamine degradation to glutamate** | | | | |
| glutaminase (EC 3.5.1.2) / glutamate synthase (NADH) (EC 1.4.1.13) | 1_20289; K337_v1_11291; K337_v1 | 60010; PSYM_v1_60010; PSY | 50; FUSI_v1_430103; FU | SP_v1_100037; DESP_v1_100 |

| | | | | | |
|---|---|---|---|---|---|
| **Histidine degradation to glutamate** | | | | | . |
| histidin ammonia lyase (EC 4.3.1.3) | K337_v1_20957 | PSYM_v1_80008 | FUSI_v1_460027 | - | . |
| urocanate hydratase (EC 4.2.1.49) | K337_v1_20351 | - | FUSI_v1_640043 | - | . |
| imidazolonepropionase (EC 3.5.2.7) | K337_v1_20956 | - | )040; FUSI_v1_680002; FUS | DESP_v1_850010 | . |
| formimidoylglutamase (EC 3.5.3.8) | K337_v1_20958 | - | - | - | . |
| **Glutamate degradation via methylaspartate pathway to pyruvate** | ~ | | | | . |
| methylaspartate mutase, mutE (EC 5.4.99.1) | K337_v1_11676; K337_v1_11678 | - | ;I_v1_940031; FUSI_v1_940 | - | . |
| methylaspartate ammonia-lyase (EC 4.3.1.2) | K337_v1_11673 | - | - | - | . |
| 2-methylmalate dehydratase (EC 4.2.1.34) | - | - | - | - | . |
| citramalate lyase (EC 4.1.3.22) | - | - | - | - | . |
| **Glutamate degradation via hydroxyglutarate pathway to butyrate** | ~ | | | | . |
| glutamate / leucine dehydrogenase (EC 1.4.1.2; EC 1.4.1.3; EC 1.4.1.4) | K337_v1_11103; K337_v1_11104 | 'M_v1_60003; PSYM_v1_291( | FUSI_v1_10079 | ˀ_v1_1130001; DESP_v1_189 | . |
| 2-hydroxyglutarate dehydrogenase (EC 1.1.99.2) | K337_v1_11115 | - | FUSI_v1_10080 | - | . |
| glutaconate CoA-transferase (EC 2.8.3.12) | - | - | - | - | . |
| 2-hydroxyglutaryl CoA dehydratase (EC 4.2.1.-) | /1_21025; K337_v1_11508; K337_v1 | - | - | - | . |
| glutaconyl-CoA decarboxylase subunit delta (EC 4.1.1.70) | K337_v1_20059 | - | - | - | . |
| glutaconyl-CoA decarboxylase subunit gamma (EC 4.1.1.70) | K337_v1_20060 | - | FUSI_v1_940037 | - | . |
| glutaconyl-CoA decarboxylase subunit beta (EC 4.1.1.70) | K337_v1_20061 | - | FUSI_v1_940038 | - | . |
| butyryl-CoA dehydrogenase (EC 1.3.8.1) | K337_v1_30044 | - | FUSI_v1_410001 | - | . |
| butyryl coenzyme A transferase, alpha subunit (EC 2.8.3.8) | K337_v1_10805 | - | - | - | . |
| butyryl coenzyme A transferase, beta subunit (EC 2.8.3.8) | K337_v1_10806 | - | - | - | . |
| **Asparagin degradation to aspartate** | | | | | . |
| asparaginase (EC 3.5.1.1) | K337_v1_20898 | M_v1_100035; PSYM_v1_100 | FUSI_v1_280010 | DESP_v1_1280014 | . |
| **L-Homocysteine degradation to cysteine and L-cysteine degradation to pyruvate** | | | | | . |
| cystathionine β-synthase (EC 4.2.1.22) | - | PSYM_v1_2510003 | I_v1_150018; FUSI_v1_111 | - | . |
| cystathionine gamma-lyase (EC 4.4.1.1) / cysteine synthase (EC 2.5.1.47) | K337_v1_11683; K337_v1_20987 | 'M_v1_660011; PSYM_v1_41( | FUSI_v1_460004 | DESP_v1_220021 | . |
| **Tryptophane degradation to pyruvate** | | | | | . |
| tryptophanase / L-cysteine desulfhydrase, PLP-dependent (EC 4.1.99.1) | K337_v1_20947 | - | - | - | . |
| **L-Serine degradation to pyruvate / Threonine degradation (I) to 2-oxobutanoate** | | | | | . |
| L-serine ammonia-lyase (EC 4.3.1.17) / threonine ammonia-lyase (EC 4.3.1.19) | K337_v1_10253; K337_v1_11287 | 'M_v1_590011; PSYM_v1_12( | )11; FUSI_v1_630018; FUS | P_v1_1300002; DESP_v1_51 | . |
| **Alanine degradation to pyruvate** | | | | | . |
| alanine dehydrogenase (EC 1.4.1.1) | K337_v1_11051; K337_v1_11052 | PSYM_v1_480006 | - | DESP_v1_20046 | . |
| **Methionine degradation to 2-oxobutanoate and methanethiol** | | | | | . |
| methionine gamma-lyase (EC 4.4.1.11) | /1_10331; K337_v1_11588; K337_v1 | PSYM_v1_20036 | )005; FUSI_v1_560010; FUS | - | . |
| **Threonine degradation (II) to glycine and acetyl-CoA** | | | | | . |
| L-threoninie 3-dehydrogenase (EC 1.1.1.103) | - | PSYM_v1_2590001 | FUSI_v1_560008 | DESP_v1_180022 | . |
| glycine C-acetyltransferase (EC 2.3.1.29) | - | PSYM_v1_2590002 | FUSI_v1_520059 | xDESP_v1_180021 | . |
| **Threonine degradation (IV) to glycine and acetyl-CoA** | | | | | . |
| low specificity L-threonine aldolase (EC 4.1.2.5) | K337_v1_11380 | - | )01; FUSI_v1_800055; FUS | ˀ_v1_2820002; DESP_v1_282 | . |
| iron-type aldehyde-alcohol dehydrogenase (NAD⁺) (EC 1.2.1.10/EC 1.1.1.1) | K337_v1_20115 | - | ;I_v1_540069; FUSI_v1_67 | )010; DESP_v1_910009; DES | . |
| **Arginine degradation (III) to putrescine** | | | | | . |
| arginine decarboxylase (EC 4.1.1.19) | K337_v1_12041 | - | FUSI_v1_1250057 | - | . |
| agmatinase (EC 3.5.3.11) | K337_v1_12038 | - | - | - | . |
| **Arginine degradation (V) to L-ornithine and CO2 (deiminase pw)** | | | | | . |
| arginine deaminase (EC 2.5.3.6) | - | - | FUSI_v1_270051 | - | . |
| ornithine carbomyltransferase (EC 2.1.3.3) | K337_v1_11897 | PSYM_v1_210021 | I_v1_270052; FUSI_v1_111 | DESP_v1_880023 | . |
| carbamate kinase (EC 2.7.2.2) | - | PSYM_v1_500009 | ;I_v1_420066; FUSI_v1_84( | - | . |
| **Lysine degradation to butyrate and acetate** | | | | | . |
| lysine 2,3-aminomutase (EC 5.4.3.2) | K337_v1_10812 | PSYM_v1_780016 | - | 20011; DESP_v1_10005; DES | . |
| lysine 5,6-aminomutase (EC 5.4.3.3) | K337_v1_10808; K337_v1_10809 | - | - | - | . |
| L-erythro-3,5-diaminohexanoate dehydrogenase (EC 1.4.1.11) | K337_v1_10813 | - | - | - | . |
| 3-keto-5-aminohexanoate cleavage enzyme | K337_v1_10814 | - | - | - | . |
| 3-aminobutyryl-CoA ammonia-lyase (EC 4.3.1.14) | K337_v1_10815 | - | - | - | . |
| acyl-CoA dehydrogenase (EC 1.3.8.1) | K337_v1_30044 | - | - | )0011; DESP_v1_1600001; DE | . |
| butyrate—acetoacetate CoA-transferase (EC 2.8.3.9) | - | - | - | - | . |
| acetyl-CoA acetyltransferase (EC 2.3.1.9) | K337_v1_30048 | - | - | )08; DESP_v1_1000008; DES | . |
| **Glycine cleavage complex** | | | | | . |
| glycine dehydrogenase (decarboxylating) (EC 1.4.4.2) | /1_20307; K337_v1_20308; K337_v1 | PSYM_v1_420003 | ;I_v1_650008; FUSI_v1_65( | )0040; DESP_v1_20041; DES | . |
| aminomethyltransferase (EC 2.1.2.10) | K337_v1_20310 | PSYM_v1_420003 | ;I_v1_650008; FUSI_v1_65( | SP_v1_20042; DESP_v1_200 | . |

| | | | | | |
|---|---|---|---|---|---|
| dihydrolipoyl dehydrogenase (EC 1.8.1.4) | K337_v1_11268; K337_v1_20310 | PSYM_v1_420003 | 006; FUSI_v1_650007; FUS | 042; DESP_v1_20045; DESP | . |
| **branched-chain α-keto acid dehydrogenase complex** | | | | | . |
| 2-keto-isovalerate dehydrogenase (EC 1.2.4.4) | - | - | FUSI_v1_810055 | - | . |
| dihydrolipoyllysine-residue (EC 2.3.1.168) | - | - | FUSI_v1_810055 | - | . |
| dihydrolipoyl dehydrogenase (EC 1.8.1.4) | - | - | _v1_1010005; FUSI_v1_11 | - | . |
| **Aminotransferases** | | | | | . |
| aminotransferase class I and II | K337_v1_10796 | - | I_v1_410019; FUSI_v1_890 | - | . |
| aminotransferase class III | - | - | FUSI_v1_640091 | - | . |
| aromatic acid aminotransferase (EC 2.6.1.57) | - | - | SI_v1_860012; FUSI_v1_10 | - | . |
| aspartate aminotransferase (EC 2.6.1.1) | K337_v1_12185 | M_v1_1700002; PSYM_v1_10 | I_v1_220004; FUSI_v1_20 | DESP_v1_120019 | . |
| branched-chain amino transferase (EC:2.6.1.42) | K337_v1_11732; K337_v1_20081 | - | I_v1_840093; FUSI_v1_620 | DESP_v1_1080001 | . |
| histidinol-phosphate aminotransferase (EC 2.6.1.9) | - | v1_3080002; PSYM_v1_1160 | FUSI_v1_360042 | DESP_v1_1210008 | . |
| alanine transaminase (EC 2.6.1.2) | - | PSYM_v1_760004 | - | - | . |
| alanine---glyoxylate transaminase (EC 2.6.1.44) | K337_v1_20164 | - | - | - | . |
| acetylornithine aminotransferase (EC 2.6.1.11) | - | PSYM_v1_80040 | SI_v1_10046; FUSI_v1_300 | DESP_v1_880024 | . |
| **Beta-oxidation of long-chain fatty acids** | | | | | . |
| Long-chain acyl-CoA synthetase (EC 6.2.1.3 / EC 6.2.1.-) (CoA-ligase) | K337_v1_10867 | M_v1_2450001; PSYM_v1_41 | - | 014; DESP_v1_450036; DES | . |
| acyl-CoA dehydrogenase (EC 1.3.8.-) | - | PSYM_v1_1300009 | FUSI_v1_410001 | 13; DESP_v1_1600001; DES | . |
| fused enoyl-CoA hydratase/isomerase ; 3-hydroxyacyl-CoA dehydrogenase (EC 4.2.1.17, EC 5.3.3.8, EC 5.1.2.3, EC 1.1.1.35) | - | PSYM_v1_3780002 | - | - | . |
| enoyl-CoA hydratase/isomerase (EC 4.2.1.17) | - | PSYM_v1_2230002 | - | 0008; DESP_v1_290019; DES | . |
| 3-hydroxyacyl-CoA dehydrogenase (EC 1.1.1.35) | - | - | - | P_v1_2710002; DESP_v1_50 | . |
| acyl-CoA thiolase (acetyl-CoA transferase) (EC 2.3.1.16) | - | M_v1_3630003; PSYM_v1_37 | - | DESP_v1_500016 | . |
| **Glycerol degradation** | | | | | . |
| glycerol-3-phosphate transporter | K337_v1_10282 | - | FUSI_v1_10096 | - | . |
| glycerol facilitator | K337_v1_10202 | - | FUSI_v1_810044 | - | . |
| glycerol kinase (sn-glycerol-3-phosphate generating) (EC 2.7.1.30) | K337_v1_10203 | - | FUSI_v1_810043 | DESP_v1_1420006 | . |
| glycerol-3-phosphate dehydrogenase (EC 1.1.5.3) | K337_v1_11183; K337_v1_12015 | - | v1_520017; FUSI_v1_96000 | DESP_v1_1240008 | . |
| **Glycolysis** | present | present | present | present | . |
| **Lactate degradation to pyruvate (reversible except for LUD-type L-lactate dehydrogenase)** | | | | | . |
| L-lactate permease, LutP and LctP | K337_v1_21046 | - | FUSI_v1_1110024 | P_v1_620009; DESP_v1_249 | . |
| L-lactate, D-lactate, and/or glycolate dehydrogenase, GlcD | K337_v1_21047 | - | I_v1_430010; FUSI_v1_111 | 0008; DESP_v1_70033; DES | . |
| L-lactate, D-lactate, and/or glycolate dehydrogenase, GlcF | - | - | - | SP_v1_620007; DESP_v1_70 | . |
| D-lactate dehydrogenase (cytochrome) (EC 1.1.2.4) | - | - | FUSI_v1_1210024; | | . |
| L-lactate dehydrogenase (EC 1.1.1.27) | - | - | FUSI_v1_460018 | - | . |
| D-lactate dehydrogenase (EC 1.1.1.28) | K337_v1_11115 | PSYM_v1_50008 | FUSI_v1_10080 | - | . |
| L-lactate dehydrogenase (EC 1.1.2.3) | - | - | - | - | . |
| LUD-type L-lactate dehydrogenase, LutABC (EC 1.1.-.-) | - | - | - | 0001, DESP_v1_450002, DES | . |
| **Butyrate degradation to butyryl-CoA (reversible)** | | | | | . |
| butyrate kinase (EC 2.7.2.7) | K337_v1_11804 | - | 0033; FUSI_v1_520026; FU | DESP_v1_480015 | . |
| phosphate butyryltransferase (EC 2.3.1.19) | K337_v1_11805 | - | 0032; FUSI_v1_520025; FU | DESP_v1_480016 | . |
| **Butyrate degradation to butyryl-CoA like in Schmidt et al., 2013 (continues with classic beta-oxidation)** | | | | | . |
| butanoate CoA-transferase (EC 2.8.3.-) | - | - | - | DESP_v1_1500004 | . |
| **Formate degradation to CO2 and H+** | | | | | . |
| formate dehydrogenase accessory protein, FdhE | - | - | - | DESP_v1_2210006 | . |
| formate dehydrogenase alpha subunit, FdhA | - | - | - | DESP_v1_2210007/8 | . |
| formate dehydrogenase beta subunit, FdhB | - | - | - | DESP_v1_2210009 | . |
| **iso-butyrate isomerization to butryl-CoA** | | | | | . |
| cob(I)alamin adenosyltransferase | K337_v1_20070 | PSYM_v1_200016 | FUSI_v1_1080006 | P_v1_1080011; DESP_v1_30 | . |
| isobutyryl-CoA mutase, N-terminal domain subunit | - | - | FUSI_v1_940031 | - | . |
| isobutyryl-CoA mutase, C-terminal domain subunit (cobalamin B12-binding domain protein) | - | - | FUSI_v1_940032 | - | . |
| LAO/AO transport system ATPase, MeaB-like protein | - | - | FUSI_v1_940033 | - | . |
| **Propionate degradation to succinyl-CoA (reversible)** | | | | - | . |
| propionyl-CoA carboxylase, gamma subunit (EC 6.4.1.3) | - | - | FUSI_v1_940035 | DESP_v1_1040006 | . |
| propionyl-CoA carboxylase, beta subunit (EC 6.4.1.3) | - | - | FUSI_v1_940037 | DESP_v1_1040008 | . |
| acetyl-CoA carboxylase, gamma subunit (EC 6.4.1.2) / propionyl-CoA carboxylase, alpha subunit (EC 6.4.1.3) | K337_v1_10646 | - | FUSI_v1_520003/4 | - | . |
| acetyl-CoA carboxylase, beta subunit (EC 6.4.1.2) / propionyl-CoA carboxylase, gamma subunit (EC 6.4.1.3) | K337_v1_10913 | - | FUSI_v1_520002 | - | . |
| methylmalonyl-CoA epimerase (EC 5.1.99.1) | - | - | FUSI_v1_940034 | - | . |
| methylmalonyl-CoA mutase accessory protein | - | - | FUSI_v1_940033 | - | . |
| methylmalonyl-CoA mutase (EC 5.4.99.2) | - | - | FUSI_v1_940031/2 | - | . |

| | | | | | |
|---|---|---|---|---|---|
| **TCA cycle** | | | ~ | | . |
| citrate synthase (EC 2.3.3.1) | - | PSYM_v1_30012 | FUSI_v1_800040 | SP_v1_90043; DESP_v1_160 | . |
| aconitate hydratase (EC 4.2.1.3) | - | M_v1_2890003; PSYM_v1_128 | FUSI_v1_800038 | 42; DESP_v1_1190001; DESI | . |
| isocitrate dehydrogenase (NAD+) (EC 1.1.1.41) / isocitrate dehydrogenase (NADP+) (EC 1.1.1.42) | - | PSYM_v1_2270005 | FUSI_v1_800039 | DESP_v1_2350002 | . |
| 2-oxoglutarate decarboxylase, thiamin-requiring (EC 1.2.4.2) [2-oxoglutarate dehydrogenase complex] | - | PSYM_v1_30006 | - | - | . |
| dihydrolipoamide succinyltransferase (EC 2.3.1.61) [2-oxoglutarate dehydrogenase complex] | - | PSYM_v1_30005 | FUSI_v1_1170017 | - | . |
| lipoamide dehydrogenase (EC 1.8.1.4) [2-oxoglutarate dehydrogenase complex] | K337_v1_11268 | PSYM_v1_420003 | 018; FUSI_v1_1010005; FU | _v1_20045; DESP_v1_86001 | . |
| succinyl-CoA synthetase, alpha chain (EC 6.2.1.5) | - | YM_v1_30002; PSYM_v1_30003 | | DESP_v1_2040003 | . |
| succinyl-CoA synthetase, beta chain (EC 6.2.1.5) | - | PSYM_v1_30004 | | DESP_v1_2040005 | . |
| succinate dehydrogenase (EC 1.3.5.1) | - | YM_v1_30008; PSYM_v1_300 | FUSI_v1_1290010 | P_v1_2040002; DESP_v1_165 | . |
| fumarate hydratase class I (EC 4.2.1.2) | K337_v1_11158/9; K337_v1_20911/2 | PSYM_v1_1700003 | _v1_1250059; FUSI_v1_12 | P_v1_1020008; DESP_v1_226 | . |
| malate dehydrogenase (EC 1.1.1.37) / Malate dehydrogenase (oxaloacetate-decarboxylating) (NADP(+)) (EC 1.1.1.40) | - | M_v1_570011; PSYM_v1_381 | - | DESP_v1_230006 | . |
| **ferrodoxin oxidoreductases** | | | | | . |
| 2-oxoglutarate ferredoxin oxidoreductase subunit alpha (EC 1.2.7.3) (KOR) | - | PSYM_v1_510008 | 029; FUSI_v1_1210029; FU | PS_v1_690022; DESP_v1_96 | . |
| 2-oxoglutarate ferredoxin oxidoreductase subunit beta (EC 1.2.7.3) (KOR) | - | PSYM_v1_510009 | 030; FUSI_v1_1210028; FU | 0004; DESP_v1_690021; DES | . |
| 2-oxoglutarate ferredoxin oxidoreductase subunit gamma (EC 1.2.7.3) (KOR) | - | - | 031; FUSI_v1_1210027; FU | DESP_v1_2130005 | . |
| 2-oxoglutarate ferredoxin oxidoreductase subunit delta (EC 1.2.7.3) (KOR) | - | - | I_v1_520028; FUSI_v1_121 | DESP_v1_2130002 | . |
| pyruvate/ketoisovalerate oxidoreductase, delta subunit, putative (EC 1.2.7.1/ EC 1.2.7.7) (POR/VOR) | K337_v1_10071 | - | SI_v1_10160; FUSI_v1_101 | DESP_v1_1080004 | . |
| pyruvate/ketoisovalerate oxidoreductase, alpha subunit (EC 1.2.7.1/ EC 1.2.7.7) (POR/VOR) | K337_v1_10072 | PSYM_v1_2910001 | SI_v1_10161; FUSI_v1_101 | DESP_v1_1080005 | . |
| pyruvate/ketoisovalerate oxidoreductase, beta subunit (EC 1.2.7.1/ EC 1.2.7.7) (POR/VOR) | K337_v1_10073 | - | SI_v1_10162; FUSI_v1_101 | DESP_v1_1080006 | . |
| pyruvate/ketoisovalerate oxidoreductase, gamma subunit(EC 1.2.7.1/ EC 1.2.7.7) (POR/VOR) | K337_v1_10074 | PSYM_v1_2910002 | SI_v1_10159; FUSI_v1_101 | DESP_v1_1080003 | . |
| tungsten-containing aldehyde:ferredoxin oxidoreductase (EC 1.2.7.5) (AOR) | K337_v1_10511 | - | FUSI_v1_1110012 | - | . |
| pyruvate-flavodoxin oxidoreductase (EC 1.2.7.-) (PFOR) | K337_v1_11862 | M_v1_2050003; PSYM_v1_253 | FUSI_v1_120078 | - | . |
| indolepyruvate:ferredoxin oxidoreductase (IOR) subunit α (EC 1.2.7.8) | - | - | FUSI_v1_800015 | DESP_v1_30060 | . |
| indolepyruvate:ferredoxin oxidoreductase (IOR) subunit β (EC 1.2.7.8) | - | - | FUSI_v1_800014 | DESP_v1_30062 | . |
| **Oxobutanoate/pyruvate degradation to propionate/acetate (via propionyl/acetyl-P intermediate)** | | | | | . |
| 2-ketobutyrate/pyruvate-formate lyase I (EC 2.3.1.54) | K337_v1_11221 | PSYM_v1_2190001 | FUSI_v1_480003 | DESP_v1_550004 | . |
| phosphate acetyltransferase (EC 2.3.1.8) | K337_v1_11864 | PSYM_v1_120027 | FUSI_v1_640081 | P_v1_960006; DESP_v1_128 | . |
| acetate kinase A and propionate kinase 2 (EC 2.7.2.1) | K337_v1_11863 | PSYM_v1_120028 | FUSI_v1_320010 | DESP_v1_1280030 | . |
| **Pyruvate dehydrogenase complex, conversion of pyruvate to acetyl-CoA** | | | | | . |
| pyruvate dehydrogenase E1 component alpha subunit (EC 1.2.4.1) | K337_v1_11265 | M_v1_420001; PSYM_v1_140 | FUSI_v1_810058 | - | . |
| pyruvate dehydrogenase E1 component beta subunit (EC 1.2.4.1) | K337_v1_11266 | PSYM_v1_2320002 | FUSI_v1_810057 | - | . |
| dihydrolipoyllysine-residue acetyltransferase (EC 2.3.1.12) | K337_v1_11267 | M_v1_2320003; PSYM_v1_42 | I_v1_810056; FUSI_v1_117 | - | . |
| dihydrolipoyl dehydrogenase (EC 1.8.1.4) | K337_v1_11268 | PSYM_v1_420003 | 018; FUSI_v1_1010005; FU | _v1_20045; DESP_v1_86001 | . |
| **Acetate production from acetyl-CoA or reverse** | ~ | | | ~ | . |
| acetyl-CoA synthetase bifunctional acetate—CoA / propionate—CoA ligase (AMP-forming) (EC 6.2.1.1/17) | - | M_v1_1400002; PSYM_v1_45 | - | - | . |
| acetate—CoA ligase (ADP-forming) (EC 6.2.1.13) | - | - | - | - | . |
| **pyruvate to oxalacetate** | | | | | . |
| pyruvate carboxylase (EC 6.4.1.1) | K337_v1_20163 | - | FUSI_v1_1200032 | - | . |
| **Sulfur cycling genes** | | | | | . |
| adenylyl-sulfate reductase, AprB (EC 1.8.99.2) | - | - | - | DESP_v1_120032 | . |
| adenylyl-sulfate reductase, AprA (EC 1.8.99.2) | - | - | - | DESP_v1_120031 | . |
| quinone-interacting membrane-bound oxidoreductase complex, QmoA | - | - | - | DESP_v1_120030 | . |
| quinone-interacting membrane-bound oxidoreductase complex, QmoB | - | - | - | DESP_v1_120029 | . |
| quinone-interacting membrane-bound oxidoreductase complex, QmoC | - | - | - | DESP_v1_120028 | . |
| sulfate adenylyltransferase, Sat (EC 2.7.7.4) | - | - | - | DESP_v1_120027 | . |
| dissimilatory sulfite reductase, DsrA (EC 1.8.99.5) | - | - | - | DESP_v1_100003 | . |
| dissimilatory sulfite reductase, DsrB (EC 1.8.99.5) | - | - | - | DESP_v1_100002 | . |
| probable regulatory protein, DsrD | - | - | - | DESP_v1_100001 | . |
| dissimilatory sulfite reductase, DsrC (EC 1.8.99.5) | - | - | - | DESP_v1_160016 | . |
| putative component of dissimilatory sulfate reduction system, DsrT | - | - | - | DESP_v1_50034 | . |
| sulfite reduction-associated membrane complex ([DsrC]-trisulfide reductase), DsrM (1.8.5.M1) | - | - | - | DESP_v1_50033 | . |
| sulfite reduction-associated membrane complex ([DsrC]-trisulfide reductase), DsrK (1.8.5.M1) | - | - | - | DESP_v1_50032 | . |
| sulfite reduction-associated membrane complex ([DsrC]-trisulfide reductase), DsrJ (1.8.5.M1) | - | - | - | DESP_v1_50031 | . |
| sulfite reduction-associated membrane complex ([DsrC]-trisulfide reductase), DsrO (1.8.5.M1) | - | - | - | DESP_v1_50030 | . |
| sulfite reduction-associated membrane complex ([DsrC]-trisulfide reductase), DsrP (1.8.5.M1) | - | - | - | DESP_v1_50029 | . |
| probable siroheme amidase, DsrN (EC 6.3.5.M1) | - | - | - | DESP_v1_1190011 | . |
| soluble inorganic pyrophosphatase (sPPase) (EC 3.6.1.1) | K337_v1_11979 | PSYM_v1_310018 | - | DESP_v1_30033 | . |
| adenylate kinase (EC 2.7.4.3) | K337_v1_30030 | PSYM_v1_440003 | FUSI_v1_940095 | DESP_v1_1090008 | . |

| | | | | | |
|---|---|---|---|---|---|
| Anaerobic sulfite reductase, subunit A (EC 1.8.1.-) | K337_v1_20489; K337_v1_20742 | - | - | - | . |
| Anaerobic sulfite reductase, subunit B (EC 1.8.1.-) | K337_v1_20488; K337_v1_20741 | - | - | - | . |
| Anaerobic sulfite reductase, subunit C (EC 1.8.1.-) | K337_v1_20487; K337_v1_20740 | - | - | - | . |
| **Nitrate reduction to ammonium (DNRA)** | | | | | . |
| perimplasmic nitrate reductase, large subunit, NapA | - | M_v1_295000; PSYM_v1_274 | - | - | . |
| periplasmic nitrate reductase, electron transfer subunit, NapB | - | PSYM_v1_2740004 | - | - | . |
| periplasmic nitrate reductase, cytochrome c-type, NapC | - | PSYM_v1_2740003 | - | - | . |
| periplasmic nitrate reductase, chaperone, NapD | - | PSYM_v1_2950002 | - | - | . |
| periplasmic nitrate reductase, ferredoxin-type protein, NapF | - | PSYM_v1_2950003 | - | - | . |
| periplasmic nitrate reductase, ferredoxin-type protein, NapG | - | PSYM_v1_2740006 | - | - | . |
| periplasmic nitrate reductase, ferredoxin-type protein, NapH | - | PSYM_v1_2740005 | - | - | . |
| formate-dependent nitrite reductase, cytochrome c552, NrfA | - | - | - | - | . |
| formate-dependent nitrite reductase, cytochrome c-type protein, NrfB | - | - | - | - | . |
| formate-dependent nitrite reductase, 4Fe4S subunit, NrfC | - | PSYM_v1_150004 | - | - | . |
| formate-dependent nitrite reductase, membrane subunit, NrfD | - | PSYM_v1_150005 | - | - | . |
| **Fumarate reduction** | | | | | . |
| fumarate reductase, flavoprotein subunit | - | PSYM_v1_1240008 | - | - | . |
| fumarate reductase iron-sulfur subunit | - | PSYM_v1_1240007 | - | - | . |
| fumarate reductase, subunit C | - | PSYM_v1_1240006 | - | - | . |
| fumarate reductase, D subunit | - | PSYM_v1_1240005 | - | - | . |
| **Oxygen respiration** | | | | | . |
| cbb3-type cytochrome c oxidase (EC 1.9.3.1) | - | ;YM_v1_70014; PSYM_v1_70( | - | - | . |
| cytochrome bd–type oxidase | - | 'M_v1_180033; PSYM_v1_18( | - | P_v1_4300221; DESP_v1_43 | . |
| cytochrome-c oxidase | - | - | - | ;P_v1_460010; DESP_v1_460 | . |
| **Type I SS** | | | | | . |
| TolC | - | PSYM_v1_3060001 | - | DESP_v1_360002 | . |
| HlyD | - | PSYM_v1_3060002 | - | - | . |
| HylB | - | PSYM_v1_3060003 | - | - | . |
| **Type II SS (general secretion pathway protein, gspD-N)** | | | | | . |
| GspC | - | PSYM_v1_50014 | - | DESP_v1_640009 | . |
| GspD | K337_v1_10274 | PSYM_v1_50015 | - | DESP_v1_640008 | . |
| GspE | - | PSYM_v1_50016 | - | DESP_v1_640007 | . |
| GspF | - | PSYM_v1_50017 | - | DESP_v1_2030004 | . |
| GspG | - | PSYM_v1_50018 | - | DESP_v1_640017 | . |
| GspH | - | PSYM_v1_50019 | - | - | . |
| GspI | - | PSYM_v1_50020 | - | DESP_v1_640015 | . |
| GspJ | - | PSYM_v1_50022 | - | DESP_v1_640014 | . |
| GspK | - | PSYM_v1_50023 | - | DESP_v1_640013 | . |
| GspL | - | PSYM_v1_50024 | - | DESP_v1_640012 | . |
| GspM | - | PSYM_v1_50025 | - | DESP_v1_640011 | . |
| GspN | - | PSYM_v1_50026 | - | DESP_v1_640010 | . |
| **Twin arginine translocation** | | | | | . |
| TatA | - | PSYM_v1_2920002 | - | DESP_v1_130035 | . |
| TatB | - | PSYM_v1_2920003 | - | DESP_v1_400009 | . |
| TatC | - | PSYM_v1_2920004 | - | DESP_v1_400010 | . |
| TatD (not needed, e.g. . Goosens et al. 2014: TAT system) | - | PSYM_v1_2920005 | - | - | . |
| **Sec pathway** | | | | | . |
| SecA | K337_v1_11974 | PSYM_v1_530006 | FUSI_v1_940046 | DESP_v1_610007 | . |
| SecB | - | PSYM_v1_30031 | - | - | . |
| SecF | K337_v1_10344 | /_v1_1420009; PSYM_v1_242 | FUSI_v1_180008 | DESP_v1_340009 | . |
| SecD | K337_v1_10343 | /_v1_1420010; PSYM_v1_242 | FUSI_v1_180009 | DESP_v1_340010 | . |
| YajC | K337_v1_12023 | PSYM_v1_2420003 | FUSI_v1_180012 | DESP_v1_340011 | . |
| SecE | K337_v1_10236 | PSYM_v1_230024 | FUSI_v1_940061 | DESP_v1_930004 | . |
| SecG | K337_v1_10735 | PSYM_v1_1410004 | FUSI_v1_690027 | DESP_v1_1070001 | . |
| SecY | K337_v1_10010 | PSYM_v1_110032 | FUSI_v1_940094 | DESP_v1_350007 | . |
| YidC | K337_v1_10153 | PSYM_v1_80033 | FUSI_v1_230014 | DESP_v1_100022 | . |
| Ffh | K337_v1_10560 | PSYM_v1_270001 | FUSI_v1_320018 | DESP_v1_30048 | . |
| FstY | K337_v1_11865 | PSYM_v1_940007 | FUSI_v1_320016 | DESP_v1_20037 | . |
| **Type IV SS** | | | | | . |

| | | | | | |
|---|---|---|---|---|---|
| VirB2 | K337_v1_20573 | - | - | - | . |
| VirB8 | K337_v1_20574 | - | - | - | . |
| VirB9 | K337_v1_20575 | - | - | - | . |
| VirB10 | K337_v1_20576 | - | - | - | . |
| VirD4 | K337_v1_20580 | - | - | - | . |
| VirB11 | K337_v1_20582 | - | - | - | . |
| VirB3 | K337_v1_20583 | - | - | - | . |
| VirB4 | K337_v1_20584 | - | - | - | . |
| VirB5 | K337_v1_20585 | - | - | - | . |
| VirB6 | K337_v1_20586 | - | - | - | . |
| VirB1 | - | - | - | - | . |
| VirB7 | - | - | - | - | . |
| **Type V SS** | | | | | . |
| Type Va | v1_10452; K337_v1_11022;K337_v1_ | YM_v1_90026;PSYM_v1_2620 | - | - | . |
| Type Vb | K337_v1_11330 | - | - | - | . |
| Type Vc | v1_11400; K337_v1_11743;K337_v1_ | - | - | - | . |

**Supplementary Table S6. Summary of additional *Psychromonas* genomes and MAGs, and key annotations for predicted secreted lipases/esterases and peptidases/proteases.**

| Genome | Genbank assembly accession no. | Sequence size (bp) | No. of contigs | GC (%) | Complete-ness | Contam-ination | Strain hetero-geneity | RAST annotation | PsortB location prediction | CDD domain (NCBI) |
|---|---|---|---|---|---|---|---|---|---|---|
| Psychromonas ossibalaenae ATCC BAA-1528 | GCA_000381745 | 5202369 | 107 | 42 | 100 | 0 | 0 | peg.1148 Phosphodiesterase/alkaline phosphatase D | Extracellular | Phosphodiesterase/alkaline phosphatase D, PhoD |
| | | | | | | | | peg.1957 Thermolabile hemolysin precursor | Extracellular | SGNH_hydrolase |
| Psychromonas aquimarina ATCC BAA-1526 | GCA_000428725 | 5534630 | 141 | 42.5 | 100 | 0 | 0 | peg.1078 Lipase precursor (EC 3.1.1.3) | Extracellular | Triacylglycerol esterase/lipase EstA |
| | | | | | | | | peg.2731 thermolabile hemolysin | Extracellular | GDSL-like Lipase/Acylhydrolase |
| | | | | | | | | peg.3736 Thermolabile hemolysin precursor | Extracellular | Phosphodiesterase/alkaline phosphatase D, PhoD |
| | | | | | | | | peg.3412 Alkaline serine protease | Extracellular | Gluzincin Peptidase family (thermolysin-like proteinases |
| | | | | | | | | peg.4597 secreted trypsin-like serine protease | Extracellular | Trypsin-like serine protease |
| Psychromonas ingrahamii 37 | GCA_000015285 | 4559598 | 1 | 40.1 | 100 | 0.15 | 60 | peg.3001 Phosphodiesterase/alkaline phosphatase D | Extracellular | Phosphodiesterase/alkaline phosphatase D, PhoD |
| | | | | | | | | peg.1920 Phospholipase/lecithinase/hemolysin | Extracellular | SGNH_hydrolase |
| Psychromonas arctica DSM 14288 | GCA_000482725 | 4745897 | 81 | 37.8 | 99.46 | 0.75 | 66.67 | peg.2566 Phosphodiesterase/alkaline phosphatase D | Extracellular | Phosphodiesterase/alkaline phosphatase D, PhoD |
| Psychromonas hadalis ATCC BAA-638 | GCA_000420245 | 3979980 | 351 | 39.1 | 100 | 0.54 | 0 | peg.25 peptidase S8 and S53, subtilisin | Extracellular | Peptidase S8 family |
| Psychromonas sp. CNPT3 | GCA_000153405 | 3052410 | 1 | 38.6 | 100 | 0 | 0 | peg.2370 Thermolabile hemolysin precursor | Extracellular | COG3240 Phospholipase |
| Psychromonas sp. CD1 | GCA_002239585 | 1688234 | 3 | 36.9 | 91.98 | 0 | 0 | peg.1338 Thermolabile hemolysin precursor | Extracellular | COG3240 Phospholipase |
| Psychromonas sp. psych-6C06 | GCA_002835465 | 3719886 | 32 | 39.3 | 100 | 0.09 | 0 | peg.1896 Vibriolysin, extracellular zinc protease | Extracellular | Gluzincin Peptidase family (thermolysin-like proteinases |
| Psychromonas sp. RZ22 | GCA_004376845 | 3069985 | 38 | 35.9 | 100 | 0 | 0 | . | . | . |
| Psychromonas sp. SP041 | GCA_000470315 | 4801754 | 121 | 36.4 | 100 | 7.21 | 28 | peg.2919 Phosphodiesterase/alkaline phosphatase D | Extracellular | Phosphodiesterase/alkaline phosphatase D, PhoD |
| Psychromonas sp. RZ5 | GCA_004378355 | 3511338 | 116 | 36.7 | 100 | 0.63 | 0 | . | . | . |
| Psychromonas sp. Urea-02u-13 | GCA_002835995 | 4749264 | 320 | 39 | 100 | 0.23 | 83.33 | peg.584 Alkaline serine protease | Extracellular | Zinc-dependent metalloprotease |
| Psychromonas sp. MB-3u-54 | GCA_002836415 | 4371403 | 186 | 40.8 | 100 | 1.02 | 28.57 | peg. 837 Phospholipase/lecithinase/hemolysin | Extracellular | SGNH_hydrolase |
| Psychromonas sp. B3M02 | GCA_003318165 | 3861573 | 156 | 38.4 | 100 | 0 | 0 | peg.969 Phosphodiesterase/alkaline phosphatase D | Extracellular | Phosphodiesterase/alkaline phosphatase D, PhoD |
| Psychromonas sp. PRT-SC03 | GCA_001321825 | 906655 | 210 | 37.1 | 30.45 | 0.86 | 66.67 | None, although incomplete genome | . | . |
| Psychromonas sp. MARI | GCA_002848885 | 1487260 | 464 | 36.9 | 84.52 | 2.16 | 40 | None, although incomplete genome | . | . |
